# Bovine Formative Embryonic Stem Cell Plasticity in Embryonic and Extraembryonic Differentiation

**DOI:** 10.1101/2024.12.28.630627

**Authors:** Yue Su, Ruifeng Zhao, Yifei Fang, Guangsheng Li, Liangliang Jin, Jiaxi Liu, Ningxiao Li, Zhen Yang, Jiaqi Zhu, Neha Mishra, Deborah Kaback, Siu Pok Yee, Chuzhao Lei, Jingyue (Ellie) Duan, Xiuchun (Cindy) Tian, Young Tang

**Author notes:** Correspondence (X.C.T.), (J.E.D), (Y.T.). These authors contributed equally (Ruifeng Zhao, Yifei Fang). These authors contributed equally (Guangsheng Li, Liangliang Jin).

## Abstract

Bovine embryonic stem cells (bESCs) can greatly enhance understanding of bovine embryonic development and applications for disease-resistance, biomedical and zoonotic pre-clinical models. However, formative bESCs with distinct morphology and complete differentiation capacity are still unreported. We document here the generation of bESCs which are pluripotent both *in vitro* and *in vivo,* and efficiently converted into neural progenitor cells (NPCs) and primordial germ cell-like cells (PGCLCs) by direct differentiation. These cells exhibited distinct metabolic features from human and mouse ESCs and early embryos. Formative bESCs contributed to a wide range of cell types within embryonic and extraembryonic tissues after aggregating with mouse and bovine embryos. The establishment of bovine formative ESCs with dual developmental plasticity represents a milestone for agricultural biotechnology and decoding the underlying mechanism of *bona fide* bovine pluripotency.

**Summary statement:** Generating bovine ESCs would greatly enhance understanding of bovine embryonic development and bioengineering of cattle for disease-resistance, biomedical and zoonotic pre-clinical models.

## Introduction

The derivation, characterization and applications of embryonic stem cells (ESCs) have advanced dramatically since their first establishment in the mouse ^1^ and human ^2^. Generating bovine ESCs (bESCs) would not only contribute to the development of new agricultural biotechnology such as cultivated meat ^3,4^, production of specialty animal products through molecular breeding and genome editing ^5–7^, but also the development of stem cell- and animal-based models to study pathogenesis and treat zoonotic diseases such as bovine spongiform encephalopathy ^8^, bovine tuberculosis ^9,10^, and *Toxoplasma gondii* infection ^11–15^.

Different states of pluripotent stem cells (PSCs) correspond to distinct developmental stages of the epiblast in the embryo. Naïve PSCs closely resemble the epiblast of a mature blastocyst ^16,17^, while primed-state cells exhibit a transcription profile akin to that of the anterior epiblast in a late-gastrula-stage embryo ^18^. An intermediate or formative state has been identified between naïve and primed states, which is capable of direct induction of primordial germ cell-like cells (PGCLCs) *in vitro* ^19–21^, whereas primed and naïve PSCs are not amenable to such induction. Both mouse naïve and formative cells demonstrate germline competence in chimeras ^19–22^, whereas mouse primed state cells show limited chimeric contribution without germline incorporation.

While abundant efforts were devoted to deriving bESCs, most cells exhibited irregular colony morphology, limited pluripotent gene expression, and failed long-term self-renewal ^23–36^. The recently reported primed-state bESCs (bpESCs) ^25,26^ also displayed irregular colony morphology and lacked chimeric capacity determination in bovine, while the bovine expanded potential stem cells (bEPSCs) ^37^ showed only extraembryonic tissue contribution of mouse-bovine interspecies chimeras with low efficiency, limited and undefined cell types in early bovine chimeric embryos. So far, bovine PSCs with defined chimeric capabilities and the ability for direct *in vitro* induction of PGCs have not been reported yet.

Human and mouse peri-implantation-stage PSCs demonstrate distinct developmental potential. Unlike mouse naive epiblast and ESCs with restricted embryonic body differentiation capacity, naive human ESCs can be directly differentiated into trophoblast stem cells *in vitro* ^38,39^, and naive human ESCs and epiblasts possess the differentiation capacity towards trophectoderm and hypoblast lineages *in vitro* and *in vivo* ^40,41^. However, the developmental plasticity near implantation stage has not been reported in species other than humans. By modulating the WNT pathway and the use of a mouse ground state ESC 3i medium ^22^, we report here the successful derivation of formative bESCs which demonstrated both embryonic and extraembryonic differentiation potentials.

## Results

### Derivation and Characterization of bESCs from Bovine *In Vitro* Blastocysts

Previously, we successfully established bovine induced pluripotent stem cells (biPSCs) ^42^ in mTeSR plus medium supplemented with H3K79 methyltransferase inhibitor iDOT1L, tankyrase inhibitor IWR1, MEK inhibitor PD0325901, glycogen synthase inhibitor CHIR99021 (2i)/LIF, and PKA agonist forskolin (TiF medium). Replacing PD0325901 with mild MEK/ERK inhibitors (PD184352 and SU5402), which supported the ground state mouse ESC ^22^ capable of germline transmission (TiF2 medium herein) resulted in thicker biPSC colonies and improved pluripotent gene expression (Figures S1*A* and S1*B*).

We then tested the applicability of TiF2 medium for bESC derivation. By culturing zona pellucida (ZP)-free bovine blastocysts in TiF2 medium on mitomycin C-treated mouse embryonic fibroblasts (MEF), we successfully derived 10 bESC lines (termed JY-bESC) from 15 *in vitro* fertilized embryos that included 3 female and 7 male lines (Figure 1*A*). JY-bESCs exhibited round, compact and dome-shaped colonies (Figure 1*B*) and were SSEA4 positive (Figure 1*C*). The doubling time of the established JY-bESC lines was 19.1±0.8 h (Figure 1*D*), faster than biPSCs ^42^. JY-bESCs maintained normal karyotype even at high passages (Figure 1*E*) and were stably propagated by single cell colonization for over 80 times. Gene-editing by using CRISPR/Cas9 in JY-bESCs was achieved with a 1.0% success rate (Figures S1*C* and S1*D*). When cultured in Matrigel, JY-bESCs formed rosette-like structure (Figure S1*E*), similar to previously reported formative state PSCs ^20,21^.

**Fig. 1.**
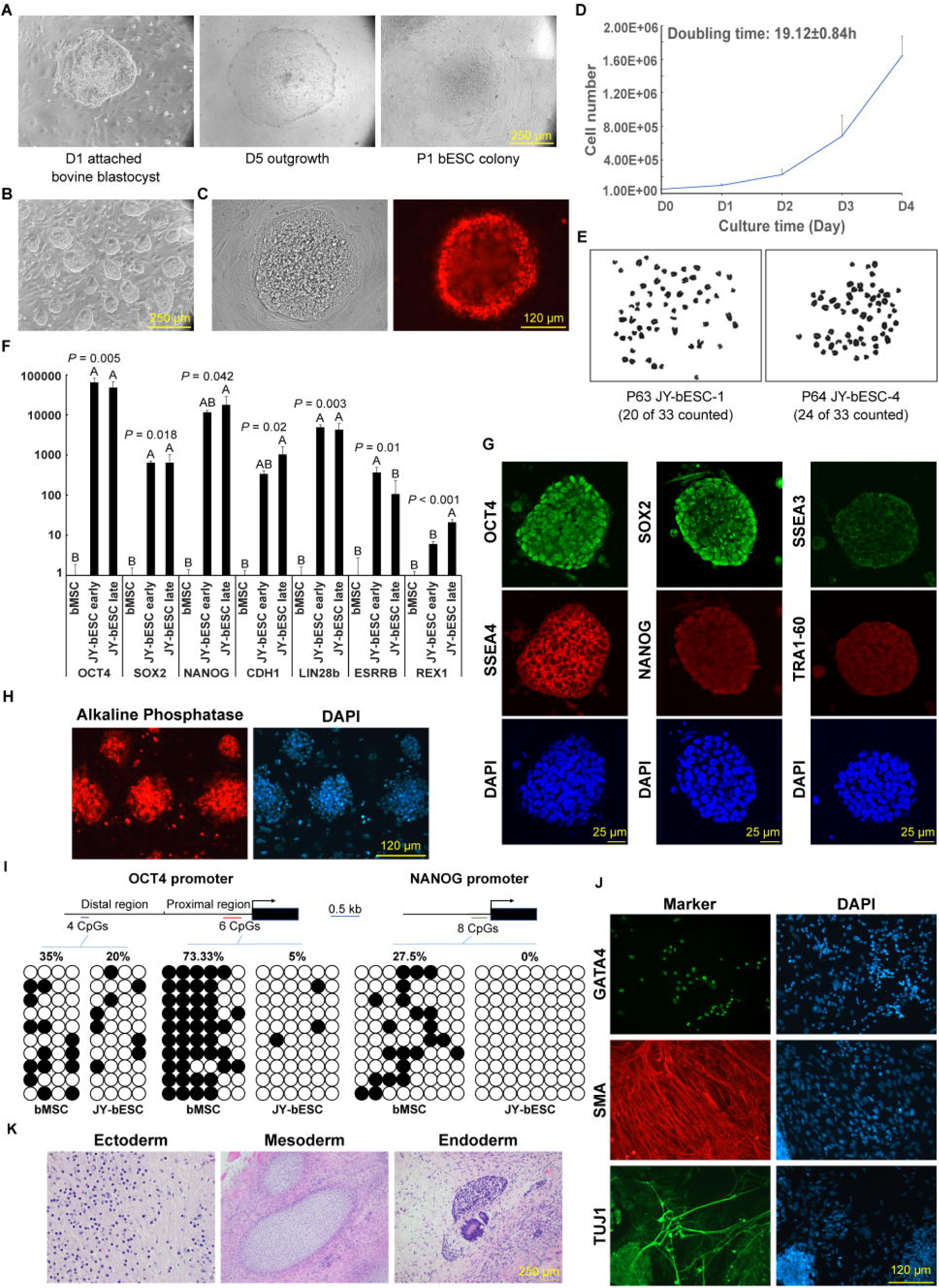
Derivation and characterization of bESCs in TiF2 medium. (*A*) Derivation of JY-bESCs cultured on feeder layers in TiF2 medium. (*B*) Colony morphology of JY-bESCs in TiF2 medium. (*C*) SSEA4 live staining of emerged colony from passaged bovine blastocyst (P2) in TiF2 medium. (*D*) Proliferation rate of JY-bESCs. Mean ± sd, n = 3, biological replicates. (*E*) Karyotypes of JY-bESC-1 and JY-bESC-4 at passages 63 and 64, respectively. (*F*) Relative pluripotent genes expression of bMSC and JY-bESCs at early (P8 to P12) and late (P42 to P50) passages. Mean ± sd, n = 3, biological replicates. (*G*) Representative immunostaining images of JY-bESCs in TiF2 medium for OCT4, NANOG, SOX2, SSEA3, SSEA4, TRA-1-60. (*H*) AP staining on JY-bESC-1 at passage 12. (*I*) Bisulfite sequencing analysis of the bovine *OCT4* distal enhancer (left two panels), proximal promoter (middle 2 panels) and *NANOG* promoter (right two panels) regions. Open and closed circles represent unmethylated and methylated CpGs, respectively. The percentage of methylated CpG is shown above each sample. (*J*) Immunostaining of passaged EBs derived from JY-bESCs for markers of the three germ layers (GATA4 for endoderm, SMA for mesoderm, and TUJ1 for ectoderm). (*K*) H&E staining of teratoma derived from JY-bESCs revealed representative tissues from the ectoderm (glial cells), mesoderm (cartilage), and endoderm (epithelial tissue with acinar complex).

JY-bESCs showed robust expression of pluripotency-related genes (*OCT4* [*a.k.a. POU5F1*], *SOX2*, *NANOG*, *E-CADHERIN* [*a.k.a. CDH1*], *LIN28B*, *ESRRB* and *REX1)* across all passages examined (Figures 1*F*, 1*G* and S1*G*). They were positive for alkaline phosphatase (AP) (Figure 1*H*) and ESC surface markers SSEA3, SSEA4 and TRA1-60 (Fig. 1*G* and Figure S1*G*). Bisulfite sequencing revealed demethylation in the *OCT4* and *NANOG* proximal promoters and partial demethylation in the distal enhancer region of *OCT4* compared to bovine mesenchymal stem cells (bMSCs) (Figure 1*I*). After the removal of growth factors, JY-bESCs successfully formed embryoid bodies (EBs) expressing markers for ectoderm, mesoderm, and endoderm (Figures 1*J*, S1*F* and S1*H*). When injected into NOD-SCID mice, JY-bESCs formed teratomas with the structures of three germ layers within four to six weeks (Figures 1*K* and S1*I*). Together, the typical ESC morphology, highly expressed pluripotency-related genes and three-germ layer differentiation both *in vitro* and *in vivo* demonstrate successful derivation of bESCs from bovine blastocysts under the TiF2 culture condition.

### Comparative Transcriptome Analysis of Bovine PSCs

To investigate the molecular signatures among different bovine PSCs, we conducted transcriptomic analysis on JY-bESCs and previously reported bEPSCs ^37^, bpESC ^26^ and biPSCs^43^ (Table S1*A*). Using principal component analysis (PCA), we found that six lines of JY-bESCs were clustered between bEPSCs and biPSCs, suggesting their distinct cellular state (Figure 2*A*). Next, we identified 643, 1,787, and 2,065 upregulated and 639, 459, and 528 downregulated genes in JY-bESCs relative to bEPSCs, biPSCs and bpESCs, respectively (Table S1*B*). Focusing on JY-bESCs, bpESCs and biPSCs, specifically, 451 genes were uniquely expressed in JY- bESCs, 275 in biPSCs, and 415 in bpESCs (Figure 2*B*). Enriched GO terms in JY-bESCs included integrin, activin, and Nodal signaling pathways (Figure 2*B*). KEGG pathway analysis highlighted ribosome biosynthesis, RAP1, and TGF-beta signaling pathways (Table S1*C*). For bpESCs, enriched GO terms and KEGG pathways included cell dedifferentiation, negative regulation of canonical WNT pathways, MAPK signaling, and FoxO pathways (Figure 2*B* and Table S1*C*). In addition, biPSCs showed enrichment in embryonic organization and fate determination GO terms, with KEGG analysis revealing Hippo signaling pathways and pluripotency regulation (Figure 2*B* and Table S1*C*). Notably, biPSCs also exhibited lower expression of *NANOG* and higher trophectoderm marker *CDX2* compared to other cells (Figures 2*B* and 2*C*).

**Fig. 2.**
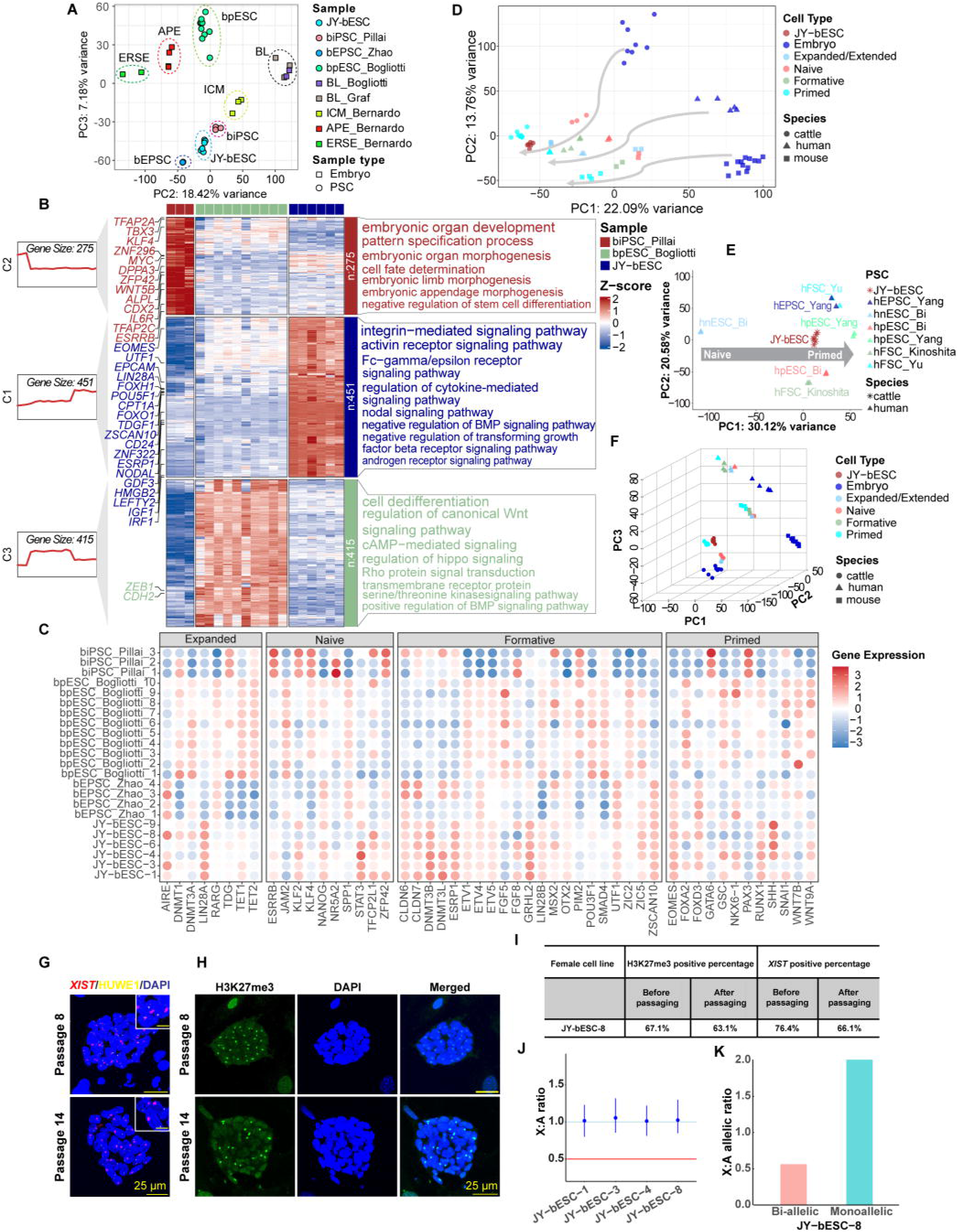
Comparative Transcriptomic and X Chromosome Inactivation (Xi) Analyses of PSCs. (*A*) PCA comparison of JY-bESCs to previously published bovine early embryos (ICM, APE, ERSE [GSE53387] and blastocyst [BL; GSE52415 and GSE110036]) and pluripotent stem cells (bpESC [GSE110036], bEPSC [GSE129760] and biPSC [GSE169624]). (*B*) Heatmap comparison of JY-bESC, biPSC_Pillai, and bpESC_Bogliotti. Key genes related to pluripotency are listed on the left. Significantly enriched Gene Ontology Biological Process (GO:BP) terms are displayed on the right. (*C*) Dot plots of marker genes representing different pluripotency states in four bovine PSCs: JY-bESC, bEPSC_Zhao, biPSC_Pillai, and bpESC_Bogliotti. (*D*) PCA plot of RNA-seq data from bovine, human, and mouse embryos and PSCs. Dot colors represent different cell types of various pluripotency states, while dot shapes indicate species. The gray arrows suggest changes in pluripotency within species. (*E*) PCA projection of JY-bESCs’ pluripotency onto that of human PSCs. (*F*) 3D PCA plot of bovine, human, and mouse embryos and PSCs. Species differences are revealed in PC3. (*G*) *XIST* and *HUWE1* RNA FISH signals and (*H*) representative H3K27me3 immunostaining images in female JY-bESCs before (top) and after (bottom) 6 additional passages. (*I*) Positive ratios of H3K27me3 and *XIST* in colonies established from single cells before and after an additional 6 passages. No significant difference (*p* > 0.05) was observed before and after the additional passages. The data were analyzed by Student’s t-test, n = 5. (*J*) X:A median expression ratios and 95% confidence intervals of four JY-bESC lines. The red line indicates failure of dosage compensation. (*K*) X:A allelic ratios of female JY-bESC-8 line.

To define the state of JY-bESCs, we analyzed their similarities to various stages of bovine embryonic development, including pre- and post-implantation embryos. Specifically, we compared PSCs with bovine dissected inner cell mass (ICM) from Day 7 blastocysts, epithelial radially symmetric epiblast (ERSE) from Day 14 embryos and anterior to posterior epiblast (APE) from Day 17 embryos ^44^. The PCA analysis revealed that the bovine PSCs were separated from the bovine embryo samples along PC1/PC2 (Figure S2*A*) and the primed state bpESCs exhibit a closer resemblance to APE while the naïve state biPSCs align closer with ICM along PC2/PC3 (Figure 2*A*). Notably, JY-bESCs were positioned intermediately between ICM and APE stages along PC2/PC3, indicating their transitional state (Figure 2*A*). We then examined known markers representing different pluripotency states ^20^ in these bovine PSCs (Figure 2*C*). These markers were defined in mice ^20^ and showed species-specific expressions across expanded, naïve, formative, and primed PSCs in mice (Figure S2*B*) and humans (Figure S2*C*). Among bovine PSCs, biPSCs had high expression of naïve pluripotency markers (*ESRRB, KLF4/2, ZFP42, NR5A2*) and upregulated primed markers (*GATA6, PAX3*) (Figure 2*C*). JY-bESCs predominantly expressed formative markers (15 out of 21) such as *CLDN6/7, DNMT3B, ESRP1, GRHL2, and LIN28B*, while also showing expression of expanded (*LIN28A*), naïve (*STAT3*, *TFCP2L1*), and primed markers (*EOMES, RUNX1*) (Figure 2*C*). Bovine primed ESCs were enriched in primed markers and showed scattered gene enrichment across other pluripotency panels (*TET1/2, JAM2, FGF5, POU3F1, FOXA2*). bEPSCs had one enriched expanded marker (*AIRE*) and elevated expression of some formative (*CLDN6, UTF1*) and primed markers (*EOMES*) (Figure 2*C*). The predominant expression of formative markers in JY-bESCs indicated a formative pluripotent status compared to other reported bovine PSCs. We observed distinct expression patterns of epigenetic modification-related genes in different bPSCs, with JY-bESCs showing enrichment in *DNMT3B*, *DNMT3L*, and *KAT7*, and lower expression of HAT1, EZH2, and KDM1A (Figure S3*A*), indicating their different states. We also compared our cells with bovine somatic cells (bovine mesenchymal stem cells, bMSCs) and found that the bMSCs exhibited enrichment in differentiation genes (*GDF6*, *SST*, *LAMA3*, *HAND1*, *DKK1*), whereas the top differentially expressed genes (DEGs) in JY-bESCs were prominently associated with pluripotency (*LIN28A*, *OTX2*, *OCT4*, *NANOG*) (Figure S3*B*). This finding confirms and emphasizes the pluripotency of JY-bESCs.

### Cross-species Analysis for Bovine-specific Features

To further identify the pluripotency status of bovine PSCs, we investigated the transcriptomic differences among PSCs of the human, mouse and bovine and their blastocysts (Table S1*A*). In general, 2D PCA (Figure 2*D*) for all three species revealed a clear cell trajectory based on pluripotency states from expanded/extended, naïve, formative, to primed PSCs. Interestingly, mouse expanded/extended PSCs (EPSC) are next to naïve PSCs, while human and bovine EPSCs are closer to formative PSCs, indicating similarity between human and bovine PSCs compared to mouse. Further projection of JY-bESC to humans or mice PSCs revealed JY-bESCs to be closest to human formative PSCs (hFSCs, Figure 2*E*), and between mouse mFSCs and primed PSCs (mpESC, Figure S4*A*).

Intriguingly, bovine, mouse, and human PSCs and blastocysts showed clear species-specific separation along PC3 of the 3D PCA (Figure 2*F*). KEGG analysis on the top 2,000 PC3 contributors revealed significant enrichment in metabolic pathways, including carbon metabolism, N-glycan biosynthesis, reactive oxygen species, purine synthesis, and biosynthesis of cofactors (Figures S5*A*-5*F*). Notably, genes that affect carbon metabolism including glycolysis (*TPI1, PGAM1, ALDOA*) and TCA cycle (*ACSS1, GLUD1, AMT, SDHA*) displayed species-specific expression patterns, with bovine showing the highest expression for *PGAM1* and *GLUD1* and comparable expression as mice for the other genes, and with humans having the lowest expression for most of these genes except for *TPI1* (Figure S5*B*). Additionally, bovine early embryonic development showed a significant shift of glycolysis intermediates towards lipid metabolism, with the lowest expression of *TPI1* (Figure S5*B*) but greatest expression of *GPD1 -* key gene for cytoplasmic glycerolipid synthesis ^45,46^ in bovine compared with humans and mice, and the lowest expression of *GPD2* which mediates the reversible interaction of *GPD1* in mitochondria (Figure S5*G*). Interestingly, *TPI1* and *GPD1* expression is much higher in bovine PSCs compared to embryos, particularly for *TPI1* in JY-bESCs (Figure S5*G*-5*H*). The distinct expression pattern of genes related to carbon metabolism indicate species-specific variations in mitochondria carbon metabolic rates among human, mouse, and bovine early embryos and PSCs.

To further define species-specific pluripotency, we compared JY-bESCs to human and mouse formative (hFSCs and mFSCs, Figure S4*B* and Table S1*D*) or primed PSCs (hpESC and mpESC, Figure S4*C* and Table S1*E*). We identified 597, 804 and 431 genes uniquely expressed in hFSCs, JY-bESCs, and mFSCs (Figure S4*B*), while 621, 606, 464 genes were specifically enriched in hpESC, JY-bESC and mpESCs, respectively (Figure S4*C*). Comparing JY-bESCs to formative PSCs revealed GO term enrichment in integrin signaling pathway for JY-bESCs and protein glycosylation and WNT signaling pathway for hFSCs and mFSCs, respectively (Figure S4*B* and Table S1*F*). Comparing JY-bESCs to primed PSCs unveiled enrichment in species-specific GO terms: extracellular matrix organization for JY-bESCs, regulation of neuron projection development for hpESCs, and ribose phosphate metabolic process for mpESCs (Figure S4*C* and Table S1*G*). Analyzing blastocysts in three species revealed 984 messages that were detected in only in the bovine embryos, with 75 genes overlapping with JY-bESC uniquely expressed genes among FSCs (Figure S4*D*). These bovine specifically expressed genes were enriched in KEGG pathways including RAP1, VEGF, and MAPK signaling pathways (*Table S*1*H*). The species-specific transcriptomic features we identified in bovine, human, and mouse PSCs and blastocysts provide insights into unique biological processes underlying early development of bovine PSC and embryos.

### Dosage Compensation of the X Chromosome in Bovine Female ESCs

Using RNA-FISH and immunostaining, we found that the female JY-bESC Line (JY-bESC-8, Figures S6*A* and S6*B*) was enriched in H3K27me3, and expressed *XIST* and a single copy of the X-linked gene *HUWE1,* suggesting X-chromosome inactivation (XaXi) (Figures 2*G* and 2*H*). After another six passages, the X chromosome status remained unchanged (Figures 2*G*, 2*H* and 2*I*). RNA-seq analysis of six JY-bESC lines (one female and five male lines) confirmed the *XIST* expression pattern (Figure S6*C*) and complete X chromosome dosage compensation (Figure 2*J*). To assess X chromosome inactivation, we further analyzed the X:A allelic ratio for bi-allelic and monoallelic single-nucleotide polymorphisms (SNPs) in the JY-bESC-8 line (Figure 2*K*). Consistent with RNA-FISH results, JY-bESC-8 showed strong monoallelic tendency (X:A allelic ratio higher than 2) and a low X:A ratio for bi-allelic genes, further supporting the establishment of XaXi status in female JY-bESCs.

### Direct Differentiation of bESCs to Neural Progenitor Cells and Primordial Germ Cell-Like Cells

The direct differentiation of bESCs such as the induction of artificial gametes and neural progenitor cells (NPCs) is critical for cell-based models to study bovine spongiform encephalopathy and perform drug screening. Also, a key feature of formative human and mouse PSCs is direct *in vitro* differentiation to PGCLCs ^19–21^. By blocking TGF-β and BMP-dependent SMAD signaling ^47^, bovine NPCs were induced within two passages (19 days), confirmed by immunostaining for NPC markers (SOX2, NESTIN, and PAX6) and elevated NPC marker gene expression (Figures 3*A-C*). Additionally, treatment with germ cell stimulator BMP4 ^48^ for four days increased expression of PGC-related genes and BMP8A further strengthened PGCLC induction (Figures 3*D* and 3*E*) as demonstrated by positive staining of PGC markers (TFAP2C, PRDM1, and SOX17) (Figures 3*F* and S1*J*). Moreover, the PRDM1+ cells exhibited co-expression with H3K27me3, while showing minimal H3K9me2 staining (Figure *3G*). This staining pattern aligns well with previous reports on PGCs ^21,49^. The triple positive ratio of TFAP2C, PRDM1 and SOX17 in bovine induced PGCLC is 21.5 ± 2.8% (Figure S1*K*), which is comparable to the rates of PGCLC induction of human and mouse formative PSCs ^19–21^. Therefore, the elevated and co-expression of NPC and PGC markers indicated efficient direct differentiation of JY-bESCs into NPC and PGCLC *in vitro*.

**Fig. 3.**
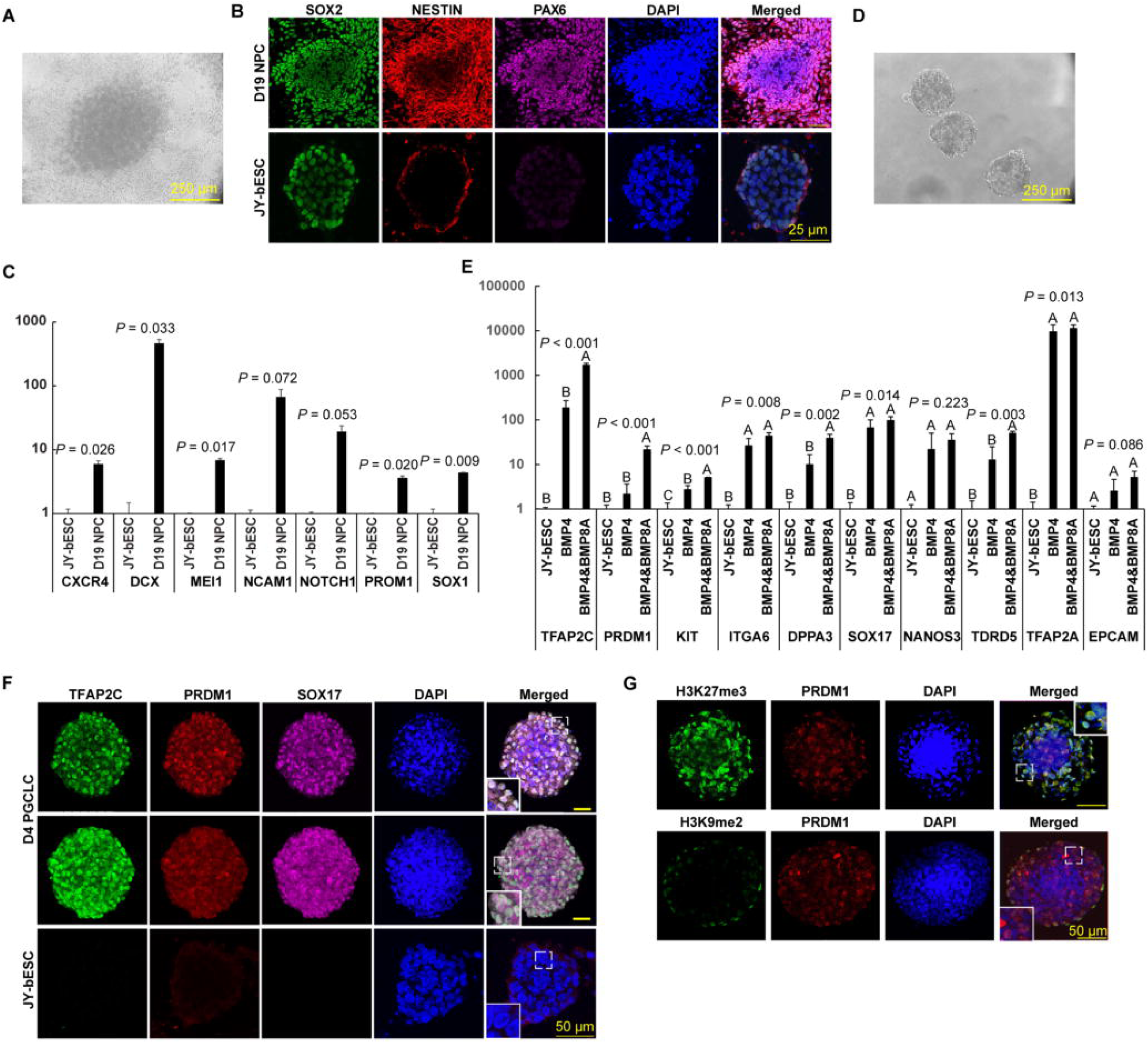
Direct *in vitro* Differentiation of bESCs to Neural Progenitor Cells (NPCs) and Primordial Germ Cell-Like Cells (PGCLCs). (*A*) The morphology of NPC 19 days after induction. (*B*) Representative immunostaining images of JY-bESCs derived NPC (upper row) and JY-bESCs (lower row) for SOX2, NESTIN, and PAX6. (*C*) Relative expression of NPC-related genes by JY-bESCs and the induced NPCs at D19. Mean ± sd, n = 2, biological replicates. (*D*) The morphology of PGCLCs four days after induction. (*E*) Relative expression of PGC-related genes by JY-bESCs and D4 PGCLC induced with BMP4, and BMP4 and BMP8a. Mean ± sd, n = 3, biological replicates. (*F*) Representative immunostaining images of PGCLCs and JY-bESCs for TFAP2C, PRDM1 and SOX17 four days after induction (first row: one layer of PGCLCs; second row: 3D projection of Z-stack scanning of PGCLCs; third row: JY-bESCs). (*G*) Representative immunostaining images of PGCLCs for H3K27me3, H3K9me2 and PRDM1 four days after induction.

### Bovine ESCs contributed to the Embryonic and Extra-embryonic Tissues of Chimeras

To determine the *in vivo* developmental potential of JY-bESCs, we established GFP-expressing (GFP+) JY-bESCs and aggregated them with mouse early morulae for the creation of interspecies chimeras ^42,50^. GFP signals were observed in the mouse blastocysts 24 hours after aggregation (Figure 4*A*) and were verified by immunofluorescence staining (Figure 4*B*), which showed that GFP+ cells were co-stained with markers of both ICM, Sox2, and trophectoderm (TE), Cdx2. Chimeric blastocysts were transferred into pseudopregnant females and retrieved on E7.5 and E8.5. Two out of eight E7.5 and three out of 44 E8.5 mouse embryos contained obvious GFP signals (Figure S7*A*). GFP signals were further detected by immunohistochemistry (IHC) with serial sections in caudal extremity, midgut, notochord, somite, and amnion of E7.5 chimeric embryos (Figures 4*C* and S7*B*). PCR analysis also confirmed the presence of bovine DNA in both embryonic and extraembryonic tissues of the remaining chimeric embryos at high frequency (33.3% embryo proper and 66.7% decidua of E7.5 and 82.8% embryo proper and 38.1% decidua of E8.5 embryos, respectively) (Figures 4*D* and S7*C*), in accordance with the blastocyst staining result. The detection of bovine cells in various tissues of mouse-bovine chimeras demonstrated the successful contribution of JY-bESCs to both embryonic and extraembryonic tissues in interspecies chimeric embryos.

**Fig. 4.**
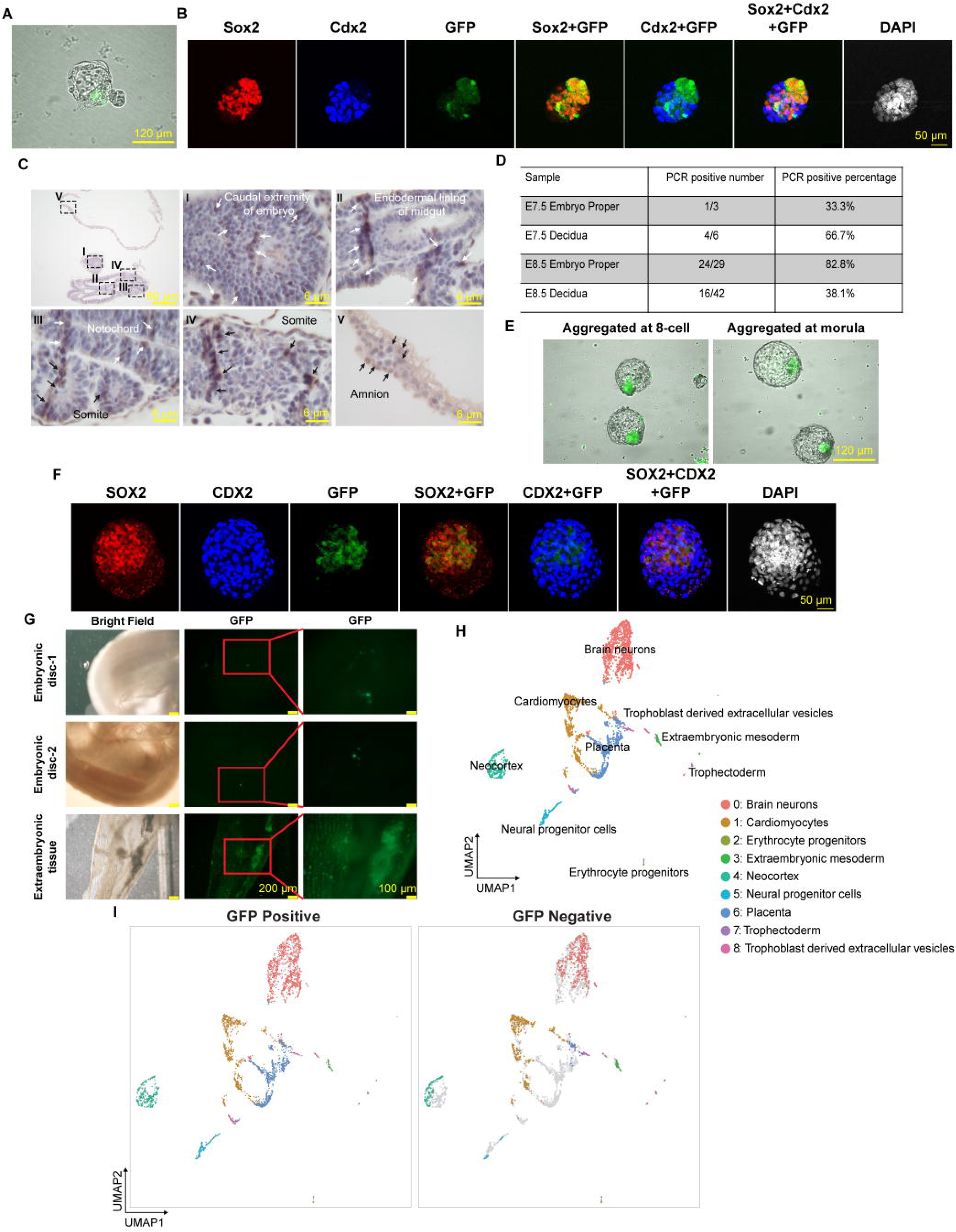
Chimeric Embryos Generated from bESCs carrying the *GFP* marker gene. (*A*) A representative mouse-bovine chimeric blastocyst 24 h after aggregation. (*B*) Whole-mount immunofluorescence of mouse-bovine chimeric blastocysts. (*C*) IHC staining of mouse-bovine E7.5 embryo using the GFP antibody. The arrows point to the positive regions. (*D*) Summary of PCR results on JY-bESCs’ contribution in mouse chimeric embryo proper and decidua. (*E*) Bovine-bovine chimeric blastocysts developed from aggregated 8-cell (left) and morula stage (right) embryos. (*F*) Whole-mount immunofluorescence of bovine chimeric. (*G*) Day 25 bovine chimeric fetus (the first and second rows) and extraembryonic tissue (the third row) with visible GFP signals. (*H*) UMAP clustering of all GFP+ cells from two D25 bovine fetuses and spiked-in cells from one extraembryonic sample. (*I*) Projection of GFP+ (GFP read count > 0) and GFP- (GFP read count = 0) cells in UMAP.

To date, extraembryonic differentiation has only been reported in bovine expanded PSCs ^37^. To this end, we aggregated GFP+ JY-bESCs with bovine embryos at the 8-cell and morula stages and further cultured them *in vitro* for 48 or 96 hours, respectively. GFP+ cells were identified in both ICM and TE of the bovine blastocysts (Figures. 4*E*, 4*F* and S7*D*). To investigate the *in vivo* development potential of the aggregated bovine chimeric embryos, 20 Day 8 GFP+ embryos were transferred to five recipient cows and five embryos from four recipients on Day 25 (Day 0 = estrus) were retrieved (Figure S7*E*). All embryos (100%) showed strong GFP signals in their extraembryonic tissues, and two out of five embryos had visible clusters of GFP+ cells in D25 fetuses by using a fluorescent microscope with 4X magnification (Figure 4*G*). FACS analysis confirmed 1.8% of the total dissociated cells from the two embryos (42,358 out of 2.2 × 10^6^) were GFP+ (Figure S7*F*).

### Single Cell Transcriptome Profiling of JY-bESC Derived Bovine Chimeras

Using 10x Genomics Chromium platform for scRNA-seq, we further examined 3,354 GFP+ JY- bESCs-derived cells and 1,083 GFP- host cells from the two Day 25 embryonic disks after spiking-in cells from one extraembryonic sample. We identified 9 clusters (See Methods for details), representing progenitors of the nerves system (Clusters 0, 4 and 5), cardiomyocytes (Cluster 1) and erythroid progenitor cells (Cluster 2) (Figures 4*H*, 4*I,* S8*A,* S8*B* and Table S2*A*). Extraembryonic mesoderm (Cluster 3) of the hypoblast origin and trophectoderm cells (Clusters 6-8) were also found, presumably from the spiked-in extraembryonic cells, which contained strong GFP signals (Figures 4*H*, 4*I,* S7*A,* S7*B* and Table S2*A*). GFP+ and GFP- cells overlapped in all the identified cell types (Figure 4*I*), indicating similar developmental trajectory by JY-bESCs and cells of the host embryos. We only identified one GFP+ cell expressing early PGC markers (*OCT4*, *KIT*, and *PRDM1*) and an additional cohort of 14 GFP+ and 4 GFP- cells that expressed at least three PGC-specific markers (*DPPA3*, *TFAP2C*, *PRDM1*, *SOX15*, *KIT*, *ITGA6*, or *ALPL*), suggesting potential PGC differentiation from our bESCs in D25 fetuses (Table S2*B* and S2*C*). Together, the diverse tissues or cell types identified in the GFP+ Day 25 bovine chimeric embryos demonstrate, for the first time, the evidence of formative bESC incorporation into both embryonic and extraembryonic tissues *in vivo*.

## Discussion

Derivation of bESCs from bovine early embryos has been challenging for decades. The previously reported bESC lines showed limited propagation ability, refractory to single-cell colonization, confined pluripotent gene expression, and restricted differentiation capacity ^23,24,27–36^. While some progress has been made in recent years with primed bESCs cultured on feeders and feeder-free substrates ^25,26^, they lacked the demonstration of chimera formation. Similarly, newly reported bovine EPSCs ^37^ showed limited interspecies chimera capability. Furthermore, unlike mouse naive PSCs, which are restricted from trophectoderm differentiation, the extraembryonic plasticity of human epiblast prior to implantation ^40,41^ has not been reported in other species. Here, we successfully derived JY-bESCs in TiF2 medium, resulting in defined colony morphology, high level expression of pluripotent genes, and *in vitro* and *in vivo* differentiation ability into three germ-layers. Notably, JY-bESCs demonstrated high rates of chimera formation to both embryonic and extraembryonic tissues in both interspecies (bovine-mouse) and intraspecies (bovine-bovine) chimeric embryos. Moreover, JY-bESCs exhibited species-specific formative transcriptomic profiles and could be directly differentiated into both NPCs and PGCLCs *in vitro*. Inter- and intraspecies chimerism further demonstrated the development potential of JY-bESCs to both embryonic and extraembryonic lineages. Taken together, JY-ESCs represent a significant advancement in extending the extraembryonic differentiation plasticity of human naive ESCs to formative bovine PSCs, offering great promises for bioengineering and development studies.

It was reported that the 3i condition (inhibited FGF receptor tyrosine kinases, ERK cascade and GSK3) maintained the ground state ESCs in mice ^22^. In our study, we used TiF2 medium containing 3i (PD184352, SU5402, and CHIR99021), and mTeSR plus with FGF2 and TGFβ, supplemented with human LIF, WNT pathway inhibitor IWR1, histone-methyltransferase inhibitor iDOT1L and the adenylcyclase activator forskolin. Different from earlier N2/B27 based 3i conditions used in two previous bESC lines ^34,35^, our TiF2 medium showed significant improvments, as evidenced by strong and homogenous pluripotent genes expression, complete differentiation in teratoma, and long-term propagation. The differences in medium components indicate that the inclusion of growth factors such as FGF2, TGFβ and LIF played crucial roles in maintaining bESC pluripotency. Notably, IWR1 has been extensively employed in recent bPSC studies ^25,26,37,42^ suggesting the essential role of blocking the WNT pathway in enhancing bESC derivation, long-term propagation and differentiation capabilities.

Our transcriptomic analysis revealed that JY-bESCs exhibit a formative state compared to other reported bovine PSCs. Cross-species analysis indicates that bovine PSCs are more similar to those from human PSCs than from mouse. Moreover, we observed varying expression of genes regulating carbon metabolism across human, mouse, and bovine early embryos and PSCs, consistent with a recent study on presomitic mesoderm cell mitochondria carbon metabolism rates ^51^. Specifically, we identified elevated *PGAM1* and *GLUD1* expression in bovine compared to humans and mice, potentially explaining species-specific differences in mitochondria respiration rates. Additionally, *TPI1* expression was highest in humans and lowest in bovine. TPI1 catalyzes conversion between dihydroxyacetone phosphate (DHAP) and glyceraldehyde-3- phosphate, which process is interconnected with glycerolipid synthesis through the conversion of DHAP to glycerol-3-phosphate by glycerol-3-phosphate dehydrogenase 1 (GPD1) ^45,46^. TPI1 deficiency was found linked to erythrocyte enzymopathy and neurological dysfunction in humans ^45,46^, suggesting human-specific glycolysis regulation for energy and neural development. These findings highlight metabolic distinctions underlying early development in different species.

Blastocyst injection is a common method for generating chimeric embryos with ESCs, but it requires expensive equipment and skilled micromanipulation. Alternatively, embryo aggregation has been used to generate chimeras ^52–54^. Here, we developed a highly relatively efficient bESC- embryo aggregation protocol, offering a convenient and effective approach for producing chimeric bovine embryos. Moreover, our scRNA-seq analysis of chimeric bovine fetuses confirmed multi-germ layer cell type differentiation from the aggregated bESCs-embryo on Day 25, known stage for bovine early PGC formation and migration ^55^, making our JY-bESCs highly valuable for *in vitro* germ cell differentiation and cattle genetic engineering.

In summary, we have successfully generated formative bESCs, which showcased a robust pluripotent gene expression profile, remarkable self-renewal potential over 80 passages with stable karyotypes, and the ability to differentiate across all three germ layers, both *in vitro* and *in vivo*. Furthermore, our JY-bESCs offer a versatile platform for precise genome editing through CRISPR-Cas9 technology and can be efficiently induced to both NPCs and PGCLCs. Notably, these cells exhibit exceptional promise for fostering interspecies and intraspecies chimerism, including their contributions to both embryonic and extraembryonic lineages. The implications of these findings are far-reaching, with the potential to catalyze innovation and propel significant advancements in the realms of agriculture and biotechnology.

### Study limitations

While successful induction of JY-bESCs to PGCLC was achieved *in vitro*, functional assays to guide their progression through meiosis and subsequent differentiation into mature gametes, either *in vitro* or *in vivo*, would be ideal to further validate their identity ^56^. Notably, our bESCs exhibited both embryonic and extraembryonic competence in chimeric embryos. Although we have demonstrated the *in vivo* development capacity of JY-bESCs into various cell types and identified putative PGCs in Day 25 fetuses, extending the gestation length, preferably to Days 35- 50 *in vivo*^57,58^, would be beneficial for the detection of oogonia and spermatogonia, in the genital ridge, as well as enable the identification of a broader array of other cell types differentiated from bESCs.

## Materials and methods

### Mice

Ten-week-old male NOD-SCID mice were used for the teratoma assay, and twelve-week-old female CD-1 mice served as recipients for the mouse-bovine chimeric embryos. The study was approved by the Institutional Animal Care and Use Committee of the University of Connecticut (protocol# A21-010) and the Health Center (protocol# 100866-0417).

### Cows

Angus cross cows, aged 2 to 6 years, were used as recipients for the bovine chimeric embryos. The study was approved by the Institutional Animal Care and Use Committee of Trans Ova Genetics (protocol# ROP68-000138).

### Bovine Embryos Production and Culture

Bovine cumulus oocyte complexes (COCs) were obtained from Applied Reproductive Technology LLC (Madison, WI). The maturation process was conducted in the HEPES-buffered IVM medium (Applied Reproductive Technology) at 38.5°C during shipping. Twenty-one to 24 hours after maturation, COCs were transferred to BO-IVF medium (IVF Bioscience) in 4-well dishes covered with mineral oil (FUJIFILM Irvine Scientific, 9305) and incubated in a 38.5°C incubator with 5.5% CO_2_. Frozen bovine semen donated by Genex was used for *in vitro* fertilization (IVF). The straws were thawed in 37°C water bath for 1-2 min and washed twice in BO-SemenPrep (IVF Bioscience) by centrifugation at 328 g for 5 min. The supernatant was discarded, and a final concentration of 1×10^6^ sperm was added (Day 0) to each IVF well at 38.5°C and 5.5% CO_2_ for 16-20 h. Presumptive zygotes were moved to 1 ml 0.76 mg/ml hyaluronidase (Sigma, H3506) for 2 min at 38.5°C to digest the remaining cumulus cells. After digestion, 800 μl supernatant was discarded, and the precipitated embryos were vortexed for 5 min to denude embryos. Denuded embryos were rinsed three times in BO-wash (IVF Bioscience) and collected to BO-IVC (IVF Bioscience) culture droplets. Embryos were incubated at 38.5°C, 5.5% CO_2_, and 6% O_2_ for further development.

### Derivation and Culture of bESCs

Bovine blastocysts produced by IVF from Days 7 to 8 were used for bESC derivation. Unhatched blastocysts were treated with freshly made 2 mg/mL Pronase (Merck Millipore, 10165921001) at 38.5°C for 2 min to remove the zona pellucida (ZP) and then thoroughly washed in IVC medium three times to remove traces of the enzyme. ZP-free blastocysts were transferred to a 24-well plate with mitomycin C treated MEF feeders in TiF2 medium that contained mTeSR-plus media (STEMCELL Technologies, 100-0276) supplemented with 2.5 μM IWR1 (AOBIOUS, AOB33702) or XAV939 (Selleck, S1180), 3.3 μM iDOT1L (AOBIOUS, AOB1922), 0.8 μM PD184352 (ApexBio, A1792), 2 μM SU5402 (ApexBio, A3843), 3 μM CHIR99021 (AOBIOUS, AOB3866), 10 μM Forskolin (AOBIOUS, AOB6380), and 1,000 U/ml human LIF (R&D Systems, 7734LF025), with the presence of 10 μM ROCKi (AOBIOUS, AOB3877) and incubated at 37°C and 5% CO_2_ for 48h without disruption. After 7-8 days, the outgrowths of the embryos were passaged using TrypLE Express (Gibco, 12604013) to a 12-well plate with feeder cells in the presence of 10 μM ROCKi. The distinguishable dome-shaped colonies started to appear from Passage 1 (P1) to P3 and were replated to fresh feeder cells with 10 μM ROCKi. The bESCs (termed JY-bESCs) were then passaged every three days at a ratio of 1:8 to 1:6 by TrypLE Express in TiF2 medium with 10 μM ROCKi.

### 3D Culture of bESCs

GFP-positive bESC lines were cultured in a 3D-system following a previously reported protocol^20^. In brief, cell pellets containing 50,000 cells were resuspended in 40 μL of ice cold Matrigel (BD Biosciences, BD356230). The solution was placed in a well of a 24-well plate and incubated at 37°C for 5 min. Then 500 μL of culture medium with 10 μM ROCKi was added to the wells. Medium was replaced every other day.

### Quantitative Reverse Transcription-PCR (qRT-PCR)

Total RNA was isolated with RNeasy mini kits (Qiagen, 74106). Genomic DNAs were removed by DNase I (Qiagen, 79254) incubation. A total of 0.5 g RNAs were then reverse transcribed into cDNA using iScript reverse transcription supermix (Bio-Rad Laboratories, 170-8891). qRT-PCR reactions were performed with SYBR Green supermix (Bimake, B21203) using the ABI 7500 Fast platform (Thermo Fisher Scientific). GAPDH was used as the housekeeping gene for expression normalization. Gene expression was determined relative to GAPDH using the ΔΔCt method with the software associated with ABI 7500. All the primers are listed in Table S3.

### Embryoid Body Differentiation

EB formation experiments were carried out with JY-bESC lines. When growing to 70–80% confluency with mainly middle-size colonies, the cells were treated with freshly prepared 1 mg/mL collagenase for 30 min and removed from the plate by pipetting. After three washes with DMEM/F12, the cells were plated onto low-adhesive petri dishes in EB formation medium (1:1 ratio of DMEM/F12 and DMEM, supplemented with 10% KSR, 5% FBS, 3 μM CHIR99021 and 20 ng/ml Activin (R&D Systems, 338AC010). Half of the medium was changed to DMEM with 10% FBS every other day. EBs were treated by 0.05% Trypsin (Gibco, 25200056) 5 days later and plated onto gelatin-coated plates. The cells were subjected to immunofluorescence staining after another 7 days of culture.

### Primordial Germ Cell Like Cell (PGCLC) Induction

Bovine PGCLCs were induced directly from JY-bESCs following a published protocol ^69^. Briefly, 3.5×10^3^ JY-bESCs were seeded into each well of ultra-low attachment round-bottom 96-well plates (Corning, 7007) in GK15 medium containing GMEM, 15% (v/v) KSR, 1× sodium pyruvate, 1× NEAA, 1× β-mercaptoethanol, 1× Glutamax and 0.5× penicillin and streptomycin, supplemented with 500 ng/mL BMP4 (GIBCO, PHC9534), 500 ng/mL BMP8a (R&D Systems, 1073BP010), 1000 U/mL LIF, 100 ng/mL SCF (R&D, 255SC010), 50 ng/mL EGF (Sigma, E9644) and 10 μM ROCKi. The cells were subjected to RNA extraction and immunostaining four days later.

### Neural Progenitor Cell (NPC) Differentiation

Bovine NPC direct differentiation protocol was modified based on the instruction of STEMdiff™ SMADi Neural Induction Kit (Stem Cell Technologies, 08581). Briefly, EBs were generated as mentioned above while the cells were cultured in STEMdiff™ Neural Induction Medium with SMAD inhibitors and 10 μM ROCKi for 4 days. The EBs were passaged to Matrigel coated wells and cultured in STEMdiff™ Neural Progenitor Medium for another 15 days when the cells were subjected to immunostaining.

### Immunostaining

The cells were first fixed in 4% paraformaldehyde (PFA) for 15 min at room temperature and then treated with 0.5% Triton X-100 in PBS for 15 min membrane permeabilization. After blocking with 5% goat serum (Cell Signaling Technology, 5425S) overnight, the cells were incubated in goat serum containing primary antibodies for 2 h at 37°C, followed by secondary antibodies at room temperature for 1 h. Similarly, the aggregated embryos at the blastocyst stage were fixed in 4% PFA for 20 min and permeabilized in 0.5% Triton X-100 for 30 min at room temperature. The embryos were blocked with goat serum and incubated with primary antibodies at 4 °C overnight and secondary antibody at room temperature for 1 h. Cells and embryos were counter-stained with DAPI and mounted on slides in Slow fade^®^ Gold antifade reagent (Invitrogen, S36938). The slides were imaged under a Nikon A1R confocal microscope.

Primary antibodies including OCT4 (Santa Crutz, sc-9081), SOX2 (Merck Millipore, AB5603), NANOG (Invitrogen, 14-5768-80), GATA4 (Cell Signaling Technology, 36966), TUJ1 (Invitrogen, A25532), SMA (Invitrogen, A25531), SOX2 (Invitrogen, 14-9811-80), NESTIN (Abcam, ab5968- 100), PAX6 (Santa Crutz, sc-81649), PRDM1 (Invitrogen, 14-5963-82), TFAP2C (Sigma, HPA055179) and SOX17 (Origene, TA500044S), H3K27me3 (Merck Millipore, 07-449), and H3K9me2 (Cell Signaling Technology, 4685s) were diluted according to the manufacturer’s instructions. Alexa Fluor 488, 594 or 647 conjugated goat anti-rabbit or goat anti-mouse secondary antibody (Cell Signaling Technology) were used in 1:1000 dilution. For cell surface marker staining, the cells were stained with NL557-conjugated TRA-1-60 (R&D Systems, NLLC4770R), SSEA-3 (R&D Systems, MAB1434) and Alexa Fluor 594 conjugated SSEA-4 (BioLegend, 330414) according to the manufacturer’s protocols.

### RNA Fluorescence in situ Hybridization (FISH)

Bovine *XIST* probes with the Quasar 570 Dye and *HUWE1* probes with the Quasar 670 Dye were purchased from Biosearch Technologies (SMF-1063-5 and SMF-1065-5). Cell fixation and hybridization were performed by following manufacture’s recommended protocols.

### Teratoma Assay

The JY-bESCs were treated with 10 μM ROCKi for 2 h before the TrypLE digestion. After digestion and centrifugation, the cells were resuspended with 30% chilly Matrigel (Corning, 354230) in DMEM/F12 and 10 μM ROCKi. 5 × 10^6^ of JY-bESCs were injected into the hint legs of 10-week-old male NOD-SCID mice (Charles River, 394NODSCID) intramuscularly (100 μl per injection). Six to eight weeks after the injection, the teratomas were dissected and fixed in 10% formalin. Paraffin-embedded teratomas were sectioned and stained with hematoxylin and eosin.

### Illumina Transcriptome Library Preparation and Sequencing

Total RNA was isolated using miRNeasy Mini kit (Qiagen, 217004). The quality of total RNA was examined with a NanoDrop 2000 spectrophotometer (Thermo Fisher Scientific, Waltham, MA, USA). To further assess RNA quality, total RNA was analyzed on the Agilent TapeStation 4200 (Agilent Technologies, Santa Clara, CA, USA) using the RNA High Sensitivity assay. Ribosomal Integrity Numbers (RINe) were recorded for each sample. Only samples with RINe values above 7.0 were considered for library preparation.

Total RNA samples were prepared for mRNA-Sequencing using the Illumina Stranded mRNA Ligation Sample Preparation kit following the manufacturer’s protocol (Illumina, San Diego, CA, USA). Libraries were validated for lengths, adapter dimers removed using the Agilent TapeStation 4200 D1000 High Sensitivity assay (Agilent Technologies, Santa Clara, CA, USA), and quantified and normalized using the dsDNA High Sensitivity Assay for Qubit 3.0 (Life Technologies, Carlsbad, CA, USA).

Sample libraries were prepared for Illumina sequencing by denaturing and diluting the libraries per manufacturer’s protocol (Illumina, San Diego, CA, USA). All samples were pooled into one sequencing pool, equally normalized, and run as one sample pool across the Illumina NovaSeq 6000 using version 1.5 chemistry. Target read depth of 50M reads was achieved per sample with paired end 100bp reads.

### Multiple Species Comparative RNA-seq Analysis

The raw FASTQ files of additional pluripotent stem cells (PSCs) were obtained from NCBI (Table S1*A*). All the data were processed with the same standard pipeline. The RNA-seq reads were mapped to the bovine genome (ARS-UCD1.2), human genome (GRCh38.p13), or mouse genome (GRCm39) obtained from Ensembl using STAR (version 2.7.10b) ^59^ with the following parameters “--quantMode TranscriptomeSAM”, “--outSAMtype BAM SortedByCoordinate”, “-- outSAMunmapped Within”, “--readFilesCommand zcat”, “--outFilterMismatchNmax 3”. The alignment data were then annotated using featureCounts ^60^ (version 2.0.3) with the following parameters: “-T 24 -p --countReadPairs -t exon -g gene_id”. For the samples sequenced using single-end reads, mapping was performed separately from the paired-end samples. To enable cross-species analysis, samples from different species were combined based on ortholog Ensembl genes.

The raw counts were used as input for the differential expression analysis, performed using the R package “DESeq2” ^70^. Genes with an adjusted P-value < 0.05 and an absolute log2 fold change ≥ 1.5 were defined as differentially expressed genes (DEGs). Principal Component Analysis (PCA) was performed using the prcomp function in R, utilizing the vst transformed values obtained from DESeq2. The PCA projection was conducted using the predict function in R. Heatmaps displaying the differentially expressed genes (DEGs) and enriched Gene Ontology Biological Process (GO: BP) terms were generated using the ’ClusterGVis’ package ^71^ in R. Additionally, the heatmap showing the expression levels of stage-specific pluripotency markers was created using the ’ggplot2’ package ^72^ in R.

### Quantification of XIST and Y-linked Genes

The adaptor removal and quality control of the raw sequencing reads were carried out using fastp (v0.23.2). The bovine reference genome (ARS-UCD 1.3) was downloaded from NCBI database, and the alignment of clean reads were performed with STAR (v2.7.9a) with default parameters (-- outFilterScoreMinOverLread 0.3, --outFilterMatchNminOverLread 0.3). The raw read counts for each gene were extracted using featureCounts (v2.0.3), and the XIST expression level was normalized by transcripts per million (TPM) using TPMCalculator. To calculate Y-linked gene expression, the Y chromosome sequence from a previous assembly (Btau_5.0.1) was added to the reference genome. We quantified the expression of five Y-linked genes, including OFD1Y, USP9Y, ZRSR2Y, DDX3Y, and EIF2S3Y.

### Dosage Compensation Analysis

For calculating the X:A expression ratio, the pairwiseCI R package was used to obtain a 95% confidence interval for the ratio of the median of X to the median of A as in a previous study ^73^. It was based on 1,000 bootstrap replicates where sampling from the original data was done with replacement and stratified by the group variables. Bootstrapping was used because it was simple to apply and did not require any distribution assumptions. Only the ubiquitous expressed genes (TPM >1) across all samples were considered in the analysis.

### SNP Calling and Allele-specific Expression Analysis

For allele-specific expression analysis, we used Varscan software to perform SNP calling (a minimum coverage of 1 and a minimum variant frequency of 0.01) with the RNA-seq data. The SNPs were further filtered by Exon region and a total number of supporting reads >=10. The ASEReadCounter module of GATK software was used to assign the mapped reads to reference allele and alternative allele. The allelic expression pattern was defined based on the fraction of reference allele expression for each SNP: if the fraction of reference allele was larger than 15% and lower than 85%, it would be defined as biallelically expressed SNP; otherwise, it would be defined as monoallelically expressed SNP.

### X:A Allelic Ratio

To determine the XCI status in the samples, we generated the X: Autosome (X:A) allelic ratio. We first counted the number of biallelic and monoallelic SNPs (which passed all filters as motioned above) on each chromosome. Since our analysis is based on expressed SNPs, this sum is expected to be correlated with the number of genes on the chromosome. To avoid this bias, we calculated a normalized SNP sum by dividing the SNP sum with the number of known genes on each chromosome. Then we calculated the ratio of normalized SNP sum of X chromosome and the average normalized SNP sum of autosomes:

NS (normalized sum) = Sum (biallelic/monoallelic SNPs) / No. of genes on chromosome X:A allelic ratio = X chromosome-NS / Average (Autosome-NS)

### Karyotyping

Karyotyping was carried out on different bESC lines. bESCs were first treated with 50 μg/ml Demecolcine (Sigma-Aldrich, D7385) at 37°C for 3 h, following which the cells were harvested by trypsinization. The cells were then incubated in hypotonic solution (0.56% KCl solution) for 15 min at 37°C. After washing in the fixative solution three times (methanol/glacial acetic acid 3:1), the cells were dropped onto wet and ice-cold glass slides. Giemsa (Sigma-Aldrich, G3032) at 1:20 dilution was applied onto the dried slides for staining. The nuclei were visualized with an Olympus microscope under a 100× oil objective lens.

### Bisulfite Sequencing

For bisulfite sequencing, genomic DNAs were extracted and bisulfite converted using the EpiTeck Bisulfite Kit (Qiagen, 59104). Bovine OCT4 and NANOG proximal promoter regions were amplified using PCR primers previously reported ^74^, and OCT4 distal primers were designed on MethPrimer ^75^. The sequences of the primers are shown in Table S3. PCR were performed with CloneAmp HiFi PCR Premix (Takara Bio USA, 639298) and cloned using the In-Fusion HD Cloning System (Clontech, 638910) into pIRES2-DsRed vector digested by BglII and EcoRI (New England Biolabs). Clones were picked, cultured in 6 mL LB medium with antibiotics overnight, and plasmid DNAs were extracted using a Qiaprep Miniprep Kit (Qiagen, 27106) and were subject to regular Sanger DNA sequencing. The results were then analyzed using the BiQ Analyzer ^76^.

### *In Vivo* Chimera Assay

We first generated a bESC fluorescent line using *piggyBac* based PB-CAG-eGFP and pCAG-PBase plasmids. The mouse-bovine chimeric embryos were then generated by aggregating mouse embryos with the GFP-positive bESC line as described previously ^50^. E2.5 mouse morulae were isolated from CD-1 females (Charles River). The ZP was removed by brief exposure to acidic Tyrode’s solution (Sigma-Aldrich, T1788) followed by three washes in KSOM embryo medium (Millipore, MR-101-D). ZP-free embryos were placed individually in micro-wells with KSOM embryo medium in an aggregation plate covered with light mineral oil (Fisher Scientific, 01211). Bovine ESCs were fed 2 h before aggregation. Cells were washed 2x with PBS, dislodged from the plates by brief exposure to 0.05% Trypsin (Sigma, SM-2003-C) to obtain clumps of 8 to 12 cells. Two clumps of cells were then placed with a ZP-free embryo in the micro- well and co-cultivated together in a 37 °C incubator with 6% CO_2_, 5% O_2_ and 89% N_2_. After an overnight incubation, bESC-mouse embryo aggregates that did not develop into blastocyst were discarded and 90 to 92 blastocysts from each line were then transferred into pseudopregnant females for subsequent development. Embryos were harvested on D7.5 and D8.5 in 4% PFA for analysis.

For the generation of bovine-bovine chimeric embryos, IVF was conducted as described before. ZP was removed from Day 3 (Day 0=fertilization) 8-cell embryos and Day 5 morulae by the treatment of 2 mg/mL Pronase for 2 min at 38.5 °C followed by three washes in SOF-HEPES medium. Similarly, the ZP-free embryos were individually cultured with three clumps of eight to twelve bESCs in the micro-wells at 38.5°C, 5% O_2_ and 5% CO_2_ until Day 7 in IVC medium. The aggregated bovine chimeric embryos were collected on Day 7 and 20 positive ones were shipped to TransOva Genetics. The embryos were transferred to five cows on Day 8 (Day 0=estrus) for further development. Embryos were flushed out on Day 25 for downstream analysis.

### Genotyping of the Mouse-bovine Chimeric Embryos

Genomic DNA was isolated from mouse embryos or extraembryonic tissues using the QIAamp DNA FFPE Tissue Kit (Qiagen, 56404). A total of 100 ng of genomic DNA was used for each PCR reaction with primers (Table S3) specific for mouse and bovine sequences using the TikTaq Hot Start PCR Master Mix (Monte Biotech, 560001).

### Immunohistochemistry (IHC)

For IHC analysis, E7.5 and E8.5 mouse embryos were isolated with their decidua from the uterus of pregnant mice. The embryos with obvious GFP signals were fixed in 4% PFA and sent to Yale Pathology Tissue Services in 70% ETOH for embedding, sectioning and IHC staining. In general, the embryos were embedded in paraffin and sectioned at 6 μm. Three sections per sample were made with 50 μm between each section. The sections were then treated with GFP antibody (Abcam, ab290). Chromogenic DAB was used as a substrate and the slides were counterstained with hematoxylin. Serial pictures of each slide were taken using an OLYMPUS AX70 microscope.

### Sample Preparation for Single-Cell RNA-Seq

Day 25 flushed bovine chimeric embryos were cut into small pieces and treated with 2 mg/ml papain (Worthington Biochemical, LS003119), 1 mg/ml DNase I (Roche, 10104159001) and 0.5x Glutamax in 2 ml Hibernate A without calcium (BrainBits, HACA500) at 37°C for 40 min. After digestion, 2 ml DPBS with 2% FBS was added to neutralize the enzyme. The embryos were singularized by pipetting 15 to 20 times. The dissociated cell suspension was passed through a 70 μm strainer and centrifuged at 300g for 5 min at 4°C. The cell pellet was resuspended in 1 ml cold RPMI medium with 2% FBS and subjected to cell sorting. After sorting, the cell pellet was resuspended in DPBS with 2% FBS at 800 cells/μL. The concentrated cells were then loaded for 10x single cell RNA sequencing.

### Single Cell 3’ Transcriptome Library Preparation and Sequencing

Prepared cells were processed for single cell transcriptome analysis using the Chromium Next GEM Single Cell 3’ GEM, Library & Gel Bead Kit v3.1 (10x Genomics, Pleasanton, CA, USA). Cells were diluted to the appropriate concentration per manufacturer recommendations prior to ensure proper loading. GEM reverse transcription, cDNA amplification and library generation followed manufacturer’s protocol (10x Genomics, Pleasanton, CA, USA).

Final libraries were validated for length and adapter dimer removal using the Agilent TapeStation 4200 D1000 High Sensitivity assay (Agilent Technologies, Santa Clara, CA, USA) then quantified and normalized using the dsDNA High Sensitivity Assay for Qubit 3.0 (Life Technologies, Carlsbad, CA, USA).

Sample libraries were prepared for Illumina sequencing by denaturing and diluting the libraries per manufacturer’s protocol (Illumina, San Diego, CA, USA). All samples were pooled into one sequencing pool with biased pooling based on variable cell numbers per sample, and run as one sample pool across the Illumina NovaSeq 6000 using version 1.5 chemistry. Target read depth of 50,000 transcriptome reads per cell was achieved in each sample with paired end reads.

### 10x scRNA-seq Data Analysis

The raw data was aligned with the bovine reference genome with the incorporation of GFP sequence and analyzed with the cell ranger pipeline (10x Genomics) to map gene expression results to each individual cell with the help of barcodes from the gel beads. We used R software v4.1.2 (https://www.r-project.org/) and Seurat v4.3.0 package ^66^ to read gene expression matrix from the .h5 file and performed further analysis. To ensure the accuracy and robustness of our results, we used DoubletFinder ^67^ to integrate multiple copies and remove potential doublets. Cells with less than 200 genes or more than 4,000 genes, and more than 5% mitochondrial gene expressed, as well as all genes not expressed in at least five cells, were filtered out. We also selected GFP-positive (GFP read count > 0) and negative cells (GFP read count = 0) for downstream analysis. In total, 4,437 high quality cells were obtained after applying these thresholds.

Data were normalized and scaled (eliminate variations introduced by the cell cycle using the vars.to.regress parameter of the ScaleData function), and highly variable genes computed using the SCTransform function. Subsequently, PCA linear dimensionality reduction analysis was performed, followed by Harmony v0.1.1 ^68^ that directly corrects the residues of the PCA for each sample. We manually adjusted the resolution to get the best clustering and identified the marker gene through the FindAllMarkers function with default parameters. We conducted automatic cell clustering with both cluster-based and single cell-based annotation ^77,78^ using known marker genes from CellMarker database ^79^. The results were then manually evaluated using data obtained from recent scRNA-seq studies that defined early embryo development and cell types of human 4-week embryos ^80^ and mouse E7.5-9.5 embryos ^81,82^. The identity of each cluster was further validated by the expression of key transcription factors (TFs) or marker genes of humans and mice from highly variable genes (HGVs) in each cluster, and by literature-mining on embryonic stage- and cell type-specific markers (Table S2*A*). The expression of marker genes in different cell types was visualized with the ggplot2 R package.

### CRISPR/Cas9-Mediated Gene Editing in bESCs

A CRISPR-Cas9 reporter vector containing the target sequence followed by an out-of-frame sequence of dTomato gene from a previous study was used for testing CRISPR/Cas9-mediated gene editing in bESCs ^83^. Briefly, this reporter vector was packaged into lentiviruses and introduced into the GFP-bESC line by spin infection as previously described ^42^. The Sp-Cas9 protein was expressed and purified following the published method ^84^. In general, the Sp-Cas9 construct was transformed into *Escherichia coli* strain BL21 (DE3) (New England Biolabs, MA, USA) and induced by 1 mM isopropylb-D-1-thiogalactopyranoside at optical density of 0.6–0.8. After overnight incubation at 23°C, cells were centrifuged and resuspended in xTractor Buffer containing DNase I, lysozyme solution, and Protease Inhibitor Cocktail (Takara Bio USA, Inc., CA). The supernatant of the sonicated cells was incubated with equilibrated TALON Metal Affinity Resin (Takara Bio USA, Inc., CA, USA) for 20 min on ice. The proteins were eluted from resin with the elution buffer (pH 7.0, 150 mM imidazole, 50 mM NaH2PO4, and 300 mM NaCl). The eluted proteins were concentrated and with buffer exchanged to phosphate-buffered saline (pH 7.4) using Protein Concentrator PES, 3K MWCO (Pierce Biotechnology, PA, USA). The purified Cas9 protein (50 pmol) was used to form the ribonucleoproteins (RNP) complex with sgRNA (250 pmol) in 37°C for 10 min. Then the RNP was transfected into 5×10^5^ bESCs with CRISPR-Cas9 reporter by electroporation. The fluorescence was observed 48 h after the transduction.

### Electroporation

Bovine ESCs (5×10^5^) in 180 μL TiF2 medium were mixed with 20 μL RNP (50 pmol Cas9 and 250 pmol sgRNA) or 10 μg PB-CAG-eGFP and pCAG-PBase plasmids (Addgene) in 20 μL Opti-MEM and added into 4 mm gene pulser cuvettes (total volume 200 μL). The voltage and capacitance used for electroporation were 200V and 500 μF. After electroporation, the cells were left in cuvette for 30 s and transferred into culture medium.

### Statistical Analysis

One way–ANOVA with Tukey’s multiple comparison test or Student’s t-test was used for data analysis. The figures were presented as mean ± standard deviation (sd). A *p*-value < 0.05(*) or 0.01(**) was considered statistically significant. All data were analyzed with SPSS platform.

## Acknowledgments

The authors thank Trans Ova Genetics for performing the chimeric embryo transfer and embryo flushing.

## Data and resource availability

RNA-seq and 10x scRNA-seq data were deposited in the Gene Expression Omnibus data repository under accession code: GSE240176.

## Author Contributions

Y.T., Y.S., X.C.T., and J.E.D. designed research; Y.S., R.Z., J.L., N.L., Z.Y., J.Z., N.M., D.K., S.P.Y., and Y.T. performed research; Y.S., R.Z., Y.F., G.L., L.J., C.L., J.E.D., X.C.T., and Y.T. analyzed data; and Y.S., J.E.D., X.C.T., and Y.T. wrote the paper.

## References

1. Evans, M.J., and Kaufman, M.H. (1981). Establishment in culture of pluripotential cells from mouse embryos. nature 292, 154–156.

2. Thomson, J.A., Itskovitz-Eldor, J., Shapiro, S.S., Waknitz, M.A., Swiergiel, J.J., Marshall, V.S., and Jones, J.M. (1998). Embryonic stem cell lines derived from human blastocysts. science 282, 1145–1147.

3. Hossain, M.S. (2019). Consumption of stem cell meat: An islamic perspective. IIUM Law Journal 27, 233–257.

4. Mengistie, D. (2020). Lab-growing meat production from stem cell. Journal of Nutrition & Food Sciences 3, 100015.

5. Kim, D., and Roh, S. (2021). Strategy to Establish Embryo-Derived Pluripotent Stem Cells in Cattle. International Journal of Molecular Sciences 22, 5011.

6. Yuan, Y. (2018). Capturing bovine pluripotency. Proceedings of the National Academy of Sciences 115, 1962–1963.

7. Zhao, J., Lai, L., Ji, W., and Zhou, Q. (2019). Genome editing in large animals: current status and future prospects. Natl Sci Rev 6, 402–420. 10.1093/nsr/nwz013.

8. Onodera, T., and Sakudo, A. (2020). Introduction to Current Progress in Advanced Research on Prions. Curr Issues Mol Biol 36, 63–66. 10.21775/cimb.036.063.

9. Olea-Popelka, F., Muwonge, A., Perera, A., Dean, A.S., Mumford, E., Erlacher-Vindel, E., Forcella, S., Silk, B.J., Ditiu, L., El Idrissi, A., et al. (2017). Zoonotic tuberculosis in human beings caused by Mycobacterium bovis-a call for action. Lancet Infect Dis 17, e21–e25. 10.1016/S1473-3099(16)30139-6.

10. Borham, M., Oreiby, A., El-Gedawy, A., Hegazy, Y., Khalifa, H.O., Al-Gaabary, M., and Matsumoto, T. (2022). Review on Bovine Tuberculosis: An Emerging Disease Associated with Multidrug-Resistant Mycobacterium Species. Pathogens 11. 10.3390/pathogens11070715.

11. Omonijo, A.O., Kalinda, C., and Mukaratirwa, S. (2022). Toxoplasma gondii Infections in Animals and Humans in Southern Africa: A Systematic Review and Meta-Analysis. Pathogens 11 10.3390/pathogens11020183.

12. Belluco, S., Simonato, G., Mancin, M., Pietrobelli, M., and Ricci, A. (2018). Toxoplasma gondii infection and food consumption: A systematic review and meta-analysis of case-controlled studies. Crit Rev Food Sci Nutr 58, 3085–3096. 10.1080/10408398.2017.1352563.

13. Sroka, J., Karamon, J., Wojcik-Fatla, A., Piotrowska, W., Dutkiewicz, J., Bilska-Zajac, E., Zajac, V., Kochanowski, M., Dabrowska, J., and Cencek, T. (2020). Toxoplasma gondii infection in slaughtered pigs and cattle in Poland: seroprevalence, molecular detection and characterization of parasites in meat. Parasit Vectors 13, 223 10.1186/s13071-020-04106-1.

14. de Barros, R.A.M., Torrecilhas, A.C., Marciano, M.A.M., Mazuz, M.L., Pereira-Chioccola, V.L., and Fux, B. (2022). Toxoplasmosis in Human and Animals Around the World. Diagnosis and Perspectives in the One Health Approach. Acta Trop 231, 106432. 10.1016/j.actatropica.2022.106432.

15. Rossi, G.A.M., de Freitas Costa, E., Gabriel, S., and Braga, F.R. (2022). A Systematic Review and Meta-Analysis on the Occurrence of Toxoplasmosis in Animals Slaughtered in Brazilian Abattoirs. Animals (Basel) 12 10.3390/ani12223102.

16. Boroviak, T., Loos, R., Bertone, P., Smith, A., and Nichols, J. (2014). The ability of inner-cell-mass cells to self-renew as embryonic stem cells is acquired following epiblast specification. Nature cell biology 16, 513–525.

17. Martello, G., and Smith, A. (2014). The nature of embryonic stem cells. Annual review of cell and developmental biology 30, 647–675.

18. Kojima, Y., Kaufman-Francis, K., Studdert, J.B., Steiner, K.A., Power, M.D., Loebel, D.A., Jones, V., Hor, A., de Alencastro, G., and Logan, G.J. (2014). The transcriptional and functional properties of mouse epiblast stem cells resemble the anterior primitive streak. Cell stem cell 14, 107–120.

19. Kinoshita, M., Barber, M., Mansfield, W., Cui, Y., Spindlow, D., Stirparo, G.G., Dietmann, S., Nichols, J., and Smith, A. (2021). Capture of mouse and human stem cells with features of formative pluripotency. Cell Stem Cell 28, 453–471. e458.

20. Wang, X., Xiang, Y., Yu, Y., Wang, R., Zhang, Y., Xu, Q., Sun, H., Zhao, Z.-A., Jiang, X., and Wang, X. (2021). Formative pluripotent stem cells show features of epiblast cells poised for gastrulation. Cell research 31, 526–541.

21. Yu, L., Wei, Y., Sun, H.X., Mahdi, A.K., Pinzon Arteaga, C.A., Sakurai, M., Schmitz, D.A., Zheng, C., Ballard, E.D., Li, J., et al. (2021). Derivation of Intermediate Pluripotent Stem Cells Amenable to Primordial Germ Cell Specification. Cell Stem Cell 28, 550–567 e512. 10.1016/j.stem.2020.11.003.

22. Ying, Q.-L., Wray, J., Nichols, J., Batlle-Morera, L., Doble, B., Woodgett, J., Cohen, P., and Smith, A. (2008). The ground state of embryonic stem cell self-renewal. Nature 453, 519–523.

23. Saito, S., Strelchenko, N., and Niemann, H. (1992). Bovine embryonic stem cell-like cell lines cultured over several passages. Roux’s archives of developmental biology 201, 134–141.

24. Kinoshita, M., Kobayashi, T., Planells, B., Klisch, D., Spindlow, D., Masaki, H., Bornelöv, S., Stirparo, G.G., Matsunari, H., and Uchikura, A. (2021). Pluripotent stem cells related to embryonic disc exhibit common self-renewal requirements in diverse livestock species. Development 148, dev199901.

25. Soto, D.A., Navarro, M., Zheng, C., Halstead, M.M., Zhou, C., Guiltinan, C., Wu, J., and Ross, P.J. (2021). Simplification of culture conditions and feeder-free expansion of bovine embryonic stem cells. Scientific reports 11, 1–15.

26. Bogliotti, Y.S., Wu, J., Vilarino, M., Okamura, D., Soto, D.A., Zhong, C., Sakurai, M., Sampaio, R.V., Suzuki, K., Izpisua Belmonte, J.C., and Ross, P.J. (2018). Efficient derivation of stable primed pluripotent embryonic stem cells from bovine blastocysts. Proc Natl Acad Sci U S A 115, 2090–2095. 10.1073/pnas.1716161115.

27. Stice, S.L., Strelchenko, N.S., Keefer, C.L., and Matthews, L. (1996). Pluripotent bovine embryonic cell lines direct embryonic development following nuclear transfer. Biology of reproduction 54, 100–110.

28. Cibelli, J.B., Stice, S.L., Golueke, P.J., Kane, J.J., Jerry, J., Blackwell, C., de León, F., and Robl, J.M. (1998). Trasgenic bovine chimeric offspring produced from somatic cell-derived stem-like cells. Nature biotechnology 16, 642–646.

29. Mitalipova, M., Beyhan, Z., and First, N.L. (2001). Pluripotency of bovine embryonic cell line derived from precompacting embryos. Cloning 3, 59–67.

30. Saito, S., Sawai, K., Ugai, H., Moriyasu, S., Minamihashi, A., Yamamoto, Y., Hirayama, H., Kageyama, S., Pan, J., and Murata, T. (2003). Generation of cloned calves and transgenic chimeric embryos from bovine embryonic stem-like cells. Biochemical and biophysical research communications 309, 104–113.

31. Wang, L., Duan, E., Sung, L.-y., Jeong, B.-S., Yang, X., and Tian, X.C. (2005). Generation and characterization of pluripotent stem cells from cloned bovine embryos. Biology of reproduction 73, 149–155.

32. Munoz, M., Rodriguez, A., De Frutos, C., Caamaño, J.N., Diez, C., Facal, N., and Gomez, E. (2008). Conventional pluripotency markers are unspecific for bovine embryonic-derived cell-lines. Theriogenology 69, 1159–1164.

33. Cao, S., Wang, F., Chen, Z., Liu, Z., Mei, C., Wu, H., Huang, J., Li, C., Zhou, L., and Liu, L. (2009). Isolation and culture of primary bovine embryonic stem cell colonies by a novel method. Journal of Experimental Zoology Part A: Ecological Genetics and Physiology 311, 368–376.

34. Park, S., Kim, D., Jung, Y.-G., and Roh, S. (2015). Thiazovivin, a Rho kinase inhibitor, improves stemness maintenance of embryo-derived stem-like cells under chemically defined culture conditions in cattle. Animal reproduction science 161, 47–57.

35. Kim, D., Park, S., Jung, Y.-G., and Roh, S. (2016). In vitro culture of stem-like cells derived from somatic cell nuclear transfer bovine embryos of the Korean beef cattle species, HanWoo. Reproduction, Fertility and Development 28, 1762–1780.

36. Wu, X., Song, M., Yang, X., Liu, X., Liu, K., Jiao, C., Wang, J., Bai, C., Su, G., and Liu, X. (2016). Establishment of bovine embryonic stem cells after knockdown of CDX2. Scientific reports 6, 1–12.

37. Zhao, L., Gao, X., Zheng, Y., Wang, Z., Zhao, G., Ren, J., Zhang, J., Wu, J., Wu, B., Chen, Y., et al. (2021). Establishment of bovine expanded potential stem cells. Proc Natl Acad Sci U S A 118. 10.1073/pnas.2018505118.

38. Cinkornpumin, J.K., Kwon, S.Y., Guo, Y., Hossain, I., Sirois, J., Russett, C.S., Tseng, H.- W., Okae, H., Arima, T., and Duchaine, T.F. (2020). Naive human embryonic stem cells can give rise to cells with a trophoblast-like transcriptome and methylome. Stem cell reports 15, 198–213.

39. Dong, C., Beltcheva, M., Gontarz, P., Zhang, B., Popli, P., Fischer, L.A., Khan, S.A., Park, K.-m., Yoon, E.-J., and Xing, X. (2020). Derivation of trophoblast stem cells from naïve human pluripotent stem cells. elife *9*, e52504.

40. Guo, G., Stirparo, G.G., Strawbridge, S.E., Spindlow, D., Yang, J., Clarke, J., Dattani, A., Yanagida, A., Li, M.A., and Myers, S. (2021). Human naive epiblast cells possess unrestricted lineage potential. Cell stem cell 28, 1040–1056. e1046.

41. Yanagida, A., Spindlow, D., Nichols, J., Dattani, A., Smith, A., and Guo, G. (2021). Naive stem cell blastocyst model captures human embryo lineage segregation. Cell stem cell 28, 1016–1022. e1014.

42. Su, Y., Wang, L., Fan, Z., Liu, Y., Zhu, J., Kaback, D., Oudiz, J., Patrick, T., Yee, S.P., and Tian, X. (2021). Establishment of Bovine-Induced Pluripotent Stem Cells. International journal of molecular sciences 22, 10489.

43. Pillai, V.V., Koganti, P.P., Kei, T.G., Gurung, S., Butler, W.R., and Selvaraj, V. (2021). Efficient induction and sustenance of pluripotent stem cells from bovine somatic cells. Biology open 10, bio058756.

44. Bernardo, A.S., Jouneau, A., Marks, H., Kensche, P., Kobolak, J., Freude, K., Hall, V., Feher, A., Polgar, Z., and Sartori, C. (2018). Mammalian embryo comparison identifies novel pluripotency genes associated with the naïve or primed state. Biology open 7, bio033282.

45. Orosz, F., Olah, J., and Ovadi, J. (2006). Triosephosphate isomerase deficiency: facts and doubts. IUBMB life 58, 703–715.

46. Orosz, F., Oláh, J., and Ovádi, J. (2009). Triosephosphate isomerase deficiency: new insights into an enigmatic disease. Biochimica et Biophysica Acta (BBA)-Molecular Basis of Disease 1792, 1168–1174.

47. Chambers, S.M., Fasano, C.A., Papapetrou, E.P., Tomishima, M., Sadelain, M., and Studer, L. (2009). Highly efficient neural conversion of human ES and iPS cells by dual inhibition of SMAD signaling. Nature biotechnology 27, 275–280.

48. Ohinata, Y., Ohta, H., Shigeta, M., Yamanaka, K., Wakayama, T., and Saitou, M. (2009). A signaling principle for the specification of the germ cell lineage in mice. Cell 137, 571–584.

49. Pirouz, M., Pilarski, S., and Kessel, M. (2013). A critical function of Mad2l2 in primordial germ cell development of mice. PLoS genetics 9, e1003712.

50. Behringer, R., Gertsenstein, M., Nagy, K.V., and Nagy, A. (2014). Manipulating the mouse embryo: a laboratory manual (Cold Spring Harbor Laboratory Press).

51. Lázaro, J., Costanzo, M., Sanaki-Matsumiya, M., Girardot, C., Hayashi, M., Hayashi, K., Diecke, S., Hildebrandt, T.B., Lazzari, G., and Wu, J. (2023). A stem cell zoo uncovers intracellular scaling of developmental tempo across mammals. Cell Stem Cell.

52. Eakin, G.S., and Hadjantonakis, A.-K. (2006). Production of chimeras by aggregation of embryonic stem cells with diploid or tetraploid mouse embryos. Nature protocols 1, 1145–1153.

53. Guo, J., Wu, B., Li, S., Bao, S., Zhao, L., Hu, S., Sun, W., Su, J., Dai, Y., and Li, X. (2014). Contribution of mouse embryonic stem cells and induced pluripotent stem cells to chimeras through injection and coculture of embryos. Stem cells international 2014.

54. Vajta, G., Peura, T., Holm, P., Paldi, A., Greve, T., Trounson, A., and Callesen, H. (2000). New method for culture of zona-included or zona-free embryos: The Well of the Well (WOW) system. Molecular Reproduction and Development: Incorporating Gamete Research 55, 256–264.

55. Wrobel, K.-H., and Süß, F. (1998). Identification and temporospatial distribution of bovine primordial germ cells prior to gonadal sexual differentiation. Anatomy and embryology 197, 451–467.

56. Handel, M.A., Eppig, J.J., and Schimenti, J.C. (2014). Applying “gold standards” to in-vitro-derived germ cells. Cell 157, 1257–1261.

57. Ross, D., Bowles, J., Hope, M., Lehnert, S., and Koopman, P. (2009). Profiles of gonadal gene expression in the developing bovine embryo. Sexual development 3, 273–283.

58. Planells, B., Gómez-Redondo, I., Sánchez, J.M., McDonald, M., Cánovas, Á., Lonergan, P., and Gutiérrez-Adán, A. (2020). Gene expression profiles of bovine genital ridges during sex determination and early differentiation of the gonads. Biology of Reproduction 102, 38–52.

59. Dobin, A., Davis, C.A., Schlesinger, F., Drenkow, J., Zaleski, C., Jha, S., Batut, P., Chaisson, M., and Gingeras, T.R. (2013). STAR: ultrafast universal RNA-seq aligner. Bioinformatics 29, 15–21.

60. Liao, Y., Smyth, G.K., and Shi, W. (2014). featureCounts: an efficient general purpose program for assigning sequence reads to genomic features. Bioinformatics 30, 923–930.

61. Love, M.I., Huber, W., and Anders, S. (2014). Moderated estimation of fold change and dispersion for RNA-seq data with DESeq2. Genome biology 15, 1–21.

62. Zhang, J. (2022). ClusterGVis: one-step to cluster and visualize gene expression matrix.

63. Vera Alvarez, R., Pongor, L.S., Mariño-Ramírez, L., and Landsman, D. (2019). TPMCalculator: one-step software to quantify mRNA abundance of genomic features. Bioinformatics 35, 1960–1962.

64. Koboldt, D.C., Zhang, Q., Larson, D.E., Shen, D., McLellan, M.D., Lin, L., Miller, C.A., Mardis, E.R., Ding, L., and Wilson, R.K. (2012). VarScan 2: somatic mutation and copy number alteration discovery in cancer by exome sequencing. Genome research 22, 568–576.

65. Van der Auwera, G.A., and O’Connor, B.D. (2020). Genomics in the cloud: using Docker, GATK, and WDL in Terra (O’Reilly Media).

66. Hao, Y., Hao, S., Andersen-Nissen, E., Mauck, W.M., Zheng, S., Butler, A., Lee, M.J., Wilk, A.J., Darby, C., and Zager, M. (2021). Integrated analysis of multimodal single-cell data. Cell 184, 3573–3587. e3529.

67. McGinnis, C.S., Murrow, L.M., and Gartner, Z.J. (2019). DoubletFinder: doublet detection in single-cell RNA sequencing data using artificial nearest neighbors. Cell systems 8, 329–337. e324.

68. Korsunsky, I., Millard, N., Fan, J., Slowikowski, K., Zhang, F., Wei, K., Baglaenko, Y., Brenner, M., Loh, P.-r., and Raychaudhuri, S. (2019). Fast, sensitive and accurate integration of single-cell data with Harmony. Nature methods 16, 1289–1296.

69. Hayashi, K., and Saitou, M. (2013). Generation of eggs from mouse embryonic stem cells and induced pluripotent stem cells. Nature protocols 8, 1513–1524.

70. Love, M.I., Huber, W., and Anders, S. (2014). Moderated estimation of fold change and dispersion for RNA-seq data with DESeq2. Genome biology 15, 1–21.

71. Zhang, J. (2022). ClusterGVis: One-step to Cluster and Visualize Gene Expression Matrix

72. Wickham, H. (2011). ggplot2. Wiley interdisciplinary reviews: computational statistics 3, 180–185.

73. Sangrithi, M.N., Royo, H., Mahadevaiah, S.K., Ojarikre, O., Bhaw, L., Sesay, A., Peters, A.H., Stadler, M., and Turner, J.M. (2017). Non-canonical and sexually dimorphic X dosage compensation states in the mouse and human germline. Developmental cell 40, 289–301. e283.

74. Cao, H., Yang, P., Pu, Y., Sun, X., Yin, H., Zhang, Y., Zhang, Y., Li, Y., Liu, Y., Fang, F., et al. (2012). Characterization of bovine induced pluripotent stem cells by lentiviral transduction of reprogramming factor fusion proteins. Int J Biol Sci 8, 498–511. 10.7150/ijbs.3723.

75. Li, L.-C., and Dahiya, R. (2002). MethPrimer: designing primers for methylation PCRs. Bioinformatics 18, 1427–1431.

76. Bock, C., Reither, S., Mikeska, T., Paulsen, M., Walter, J., and Lengauer, T. (2005). BiQ Analyzer: visualization and quality control for DNA methylation data from bisulfite sequencing. Bioinformatics 21, 4067–4068.

77. Aran, D., Looney, A.P., Liu, L., Wu, E., Fong, V., Hsu, A., Chak, S., Naikawadi, R.P., Wolters, P.J., and Abate, A.R. (2019). Reference-based analysis of lung single-cell sequencing reveals a transitional profibrotic macrophage. Nature immunology 20, 163–172.

78. Cortal, A., Martignetti, L., Six, E., and Rausell, A. (2021). Gene signature extraction and cell identity recognition at the single-cell level with Cell-ID. Nature biotechnology 39, 1095–1102.

79. Zhang, X., Lan, Y., Xu, J., Quan, F., Zhao, E., Deng, C., Luo, T., Xu, L., Liao, G., and Yan, M. (2019). CellMarker: a manually curated resource of cell markers in human and mouse. Nucleic acids research 47, D721–D728.

80. Xu, Y., Zhang, T., Zhou, Q., Hu, M., Qi, Y., Xue, Y., Nie, Y., Wang, L., Bao, Z., and Shi, W. (2023). A single-cell transcriptome atlas profiles early organogenesis in human embryos. Nature cell biology 25, 604–615.

81. Qiu, C., Cao, J., Martin, B.K., Li, T., Welsh, I.C., Srivatsan, S., Huang, X., Calderon, D., Noble, W.S., and Disteche, C.M. (2022). Systematic reconstruction of cellular trajectories across mouse embryogenesis. Nature genetics 54, 328–341.

82. Qiu, C., Martin, B.K., Welsh, I.C., Daza, R.M., Le, T.-M., Huang, X., Nichols, E.K., Taylor, M.L., Fulton, O., and Gomes, A.R. (2023). A single-cell transcriptional timelapse of mouse embryonic development, from gastrula to pup. bioRxiv.

83. D’Astolfo, D.S., Pagliero, R.J., Pras, A., Karthaus, W.R., Clevers, H., Prasad, V., Lebbink, R.J., Rehmann, H., and Geijsen, N. (2015). Efficient intracellular delivery of native proteins. Cell 161, 674–690.

84. Zhu, J., Su, Y., and Tang, Y. (2022). Disrupting ACE2 Dimerization Mitigates the Infection by SARS-CoV-2 Pseudovirus. Frontiers in Virology 2. 10.3389/fviro.2022.916700.

