## Supplementary figures and images for "Bovine Formative Embryonic Stem Cell Plasticity in Embryonic and Extraembryonic Differentiation"

### Suplemental figure 1

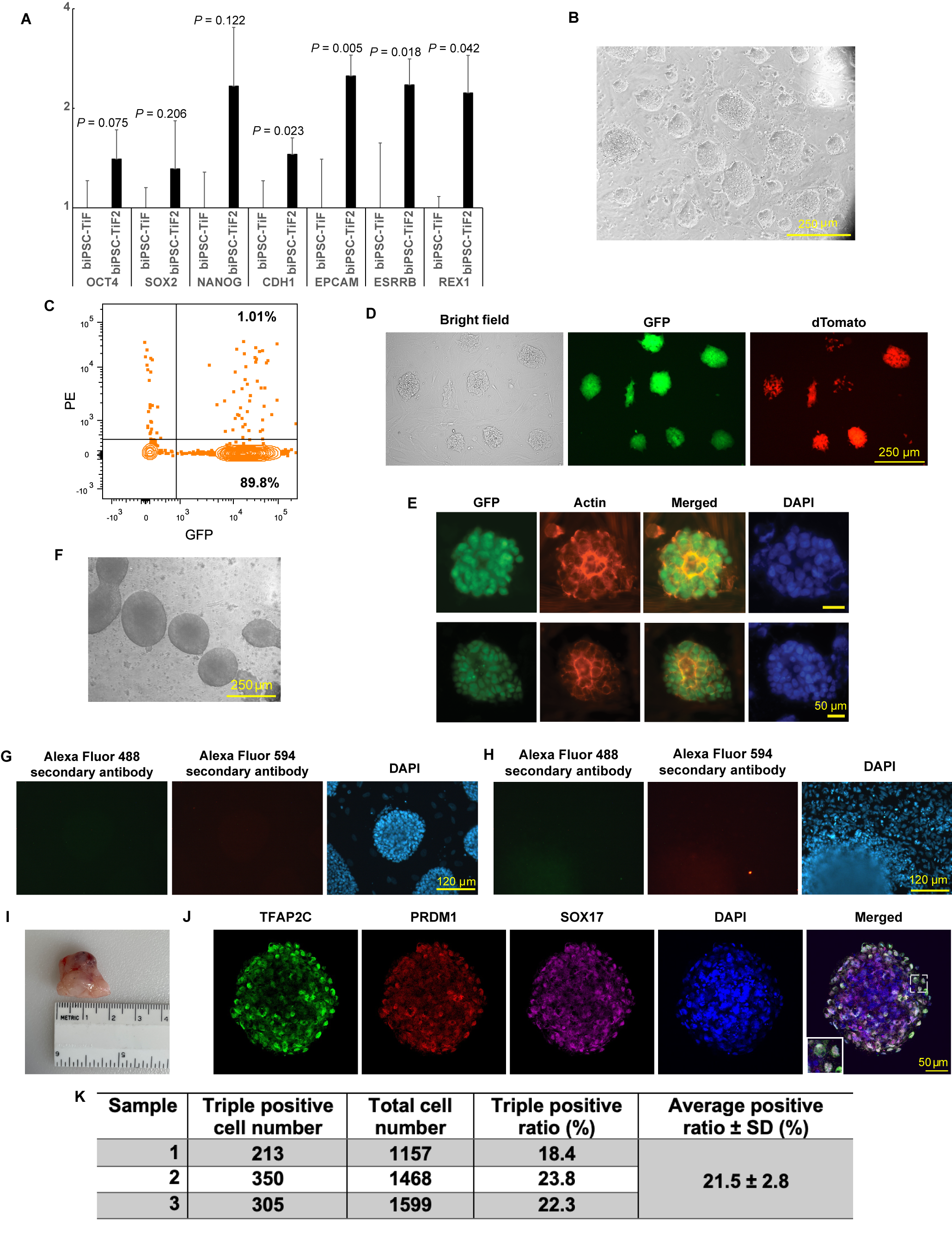

### Suplemental figure 2

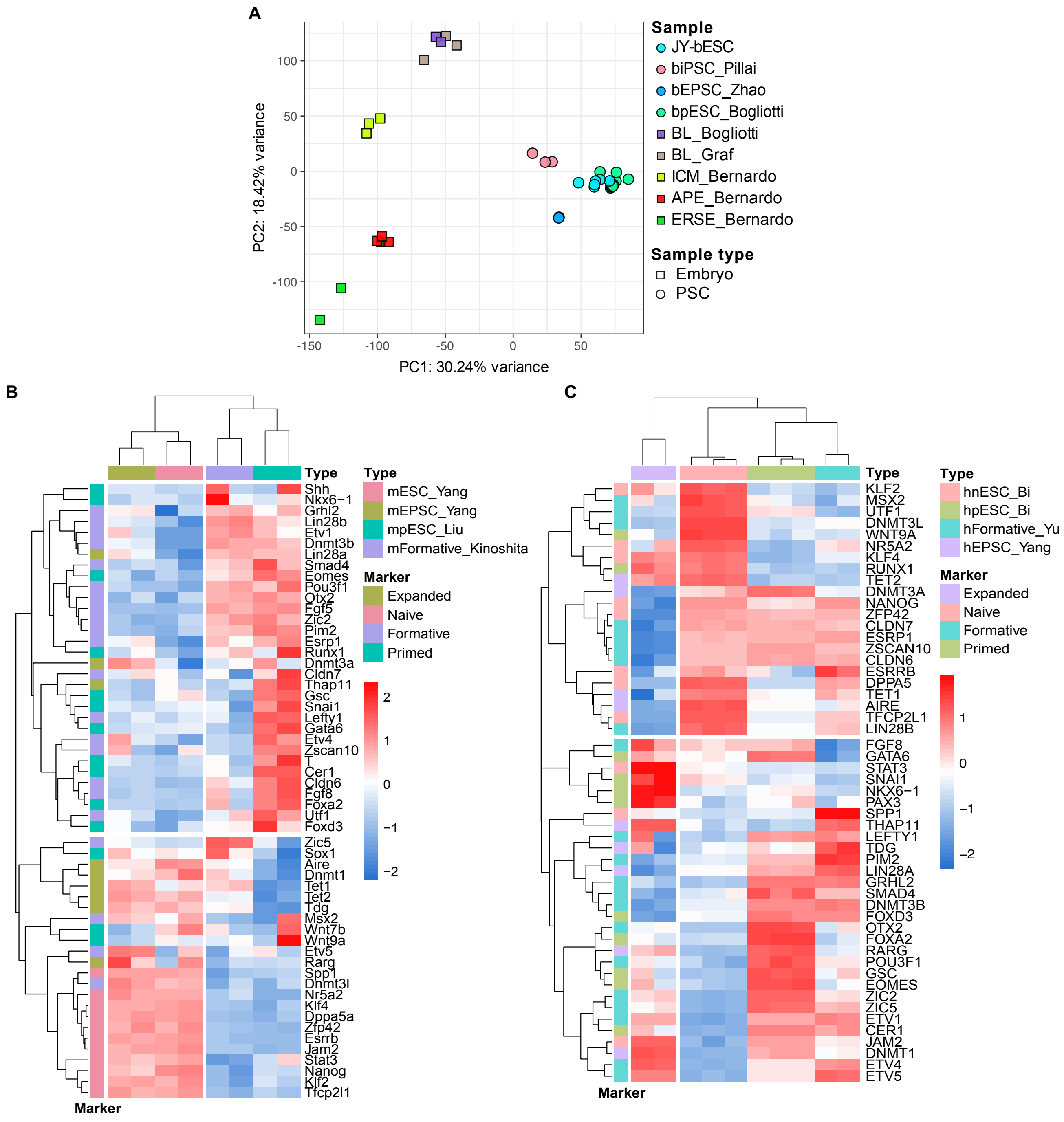

### Suplemental figure 3

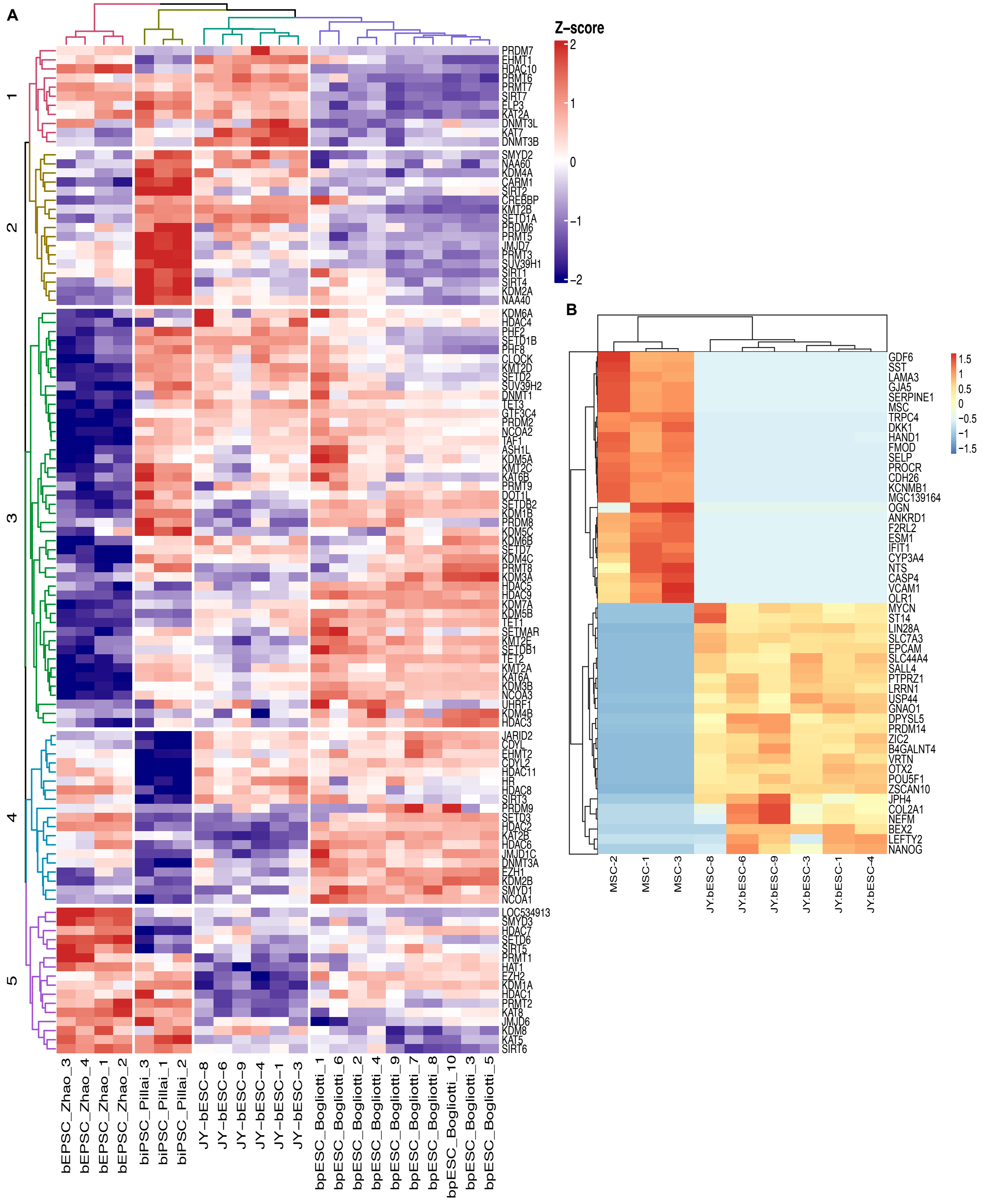

### Suplemental figure 4

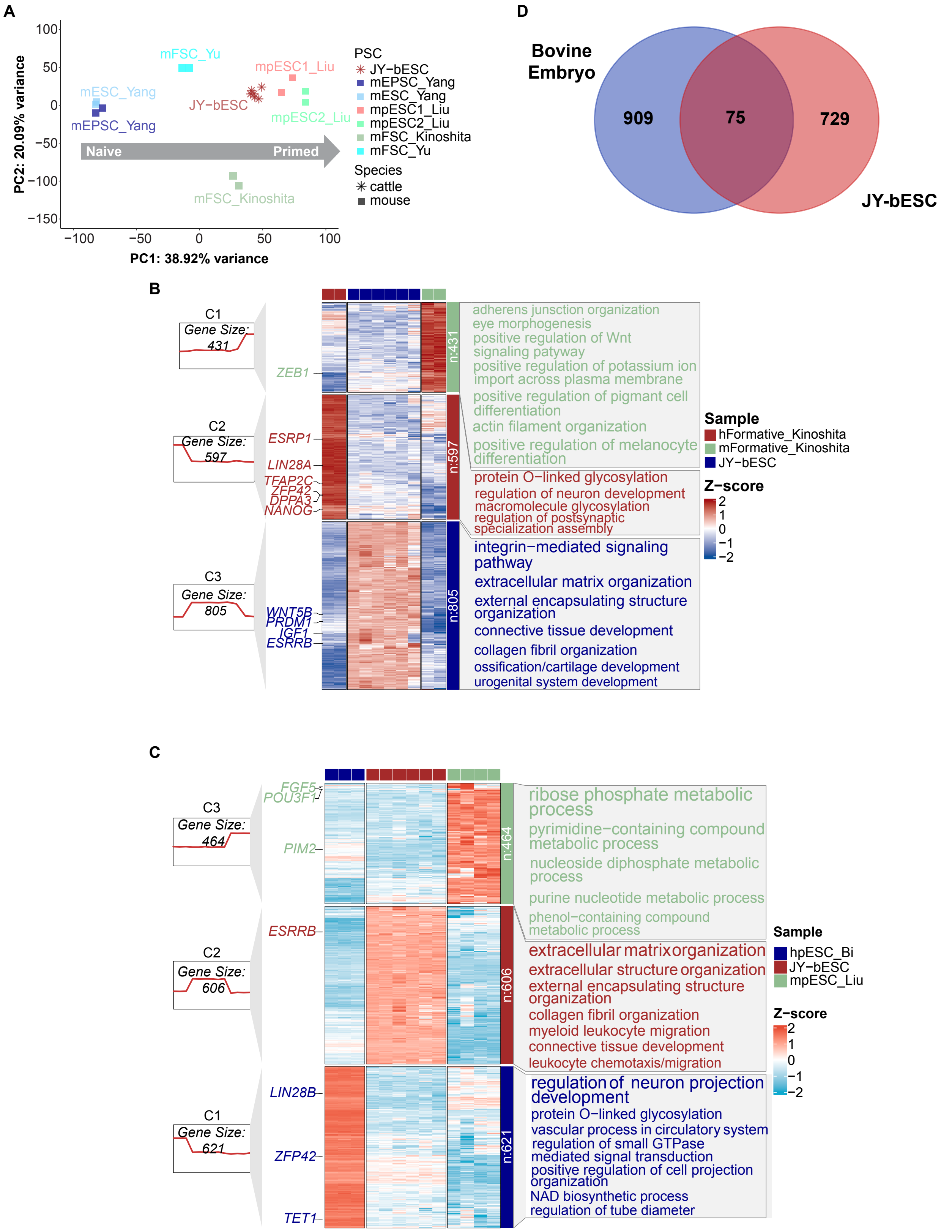

### Suplemental figure 5

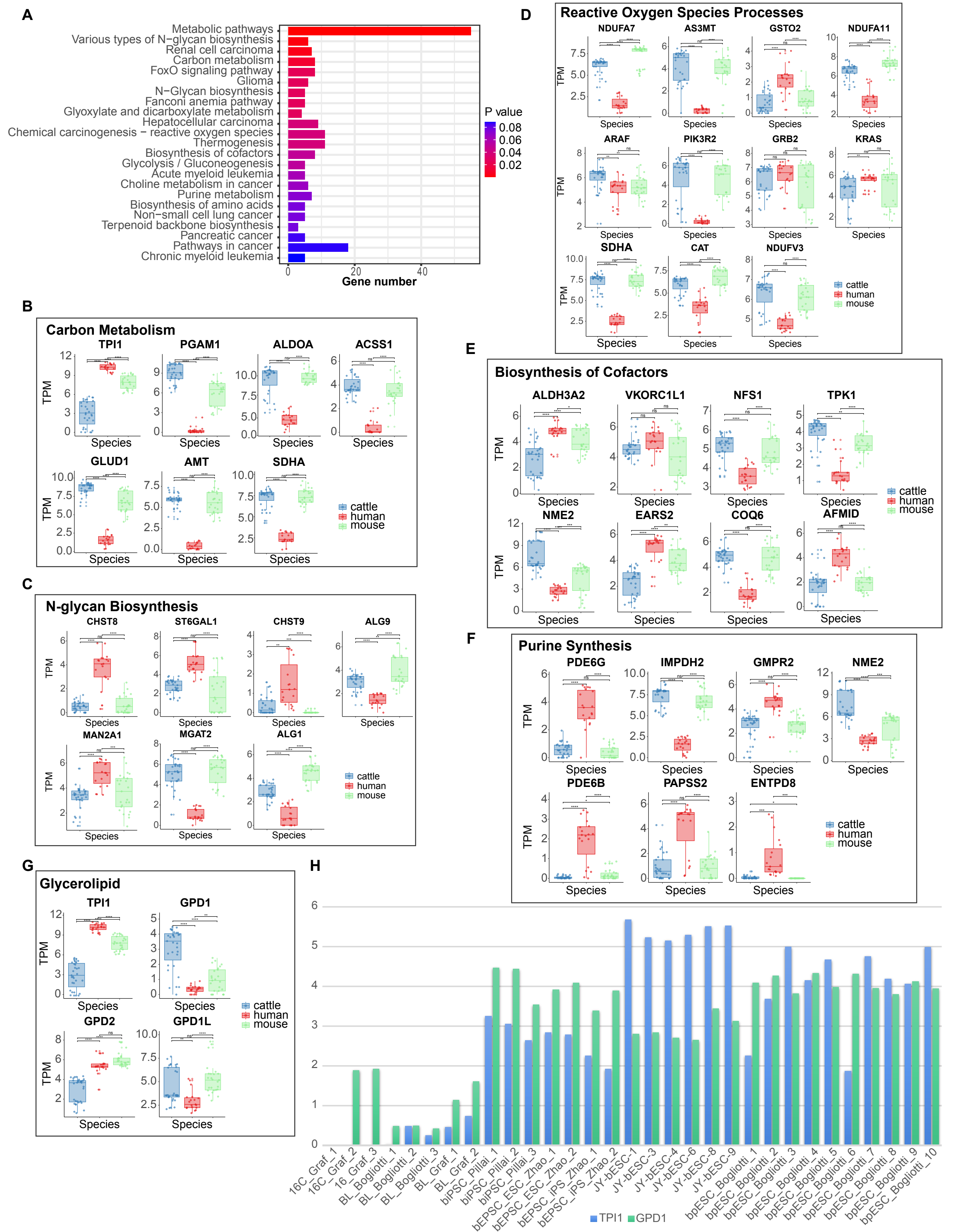

### Suplemental figure 6

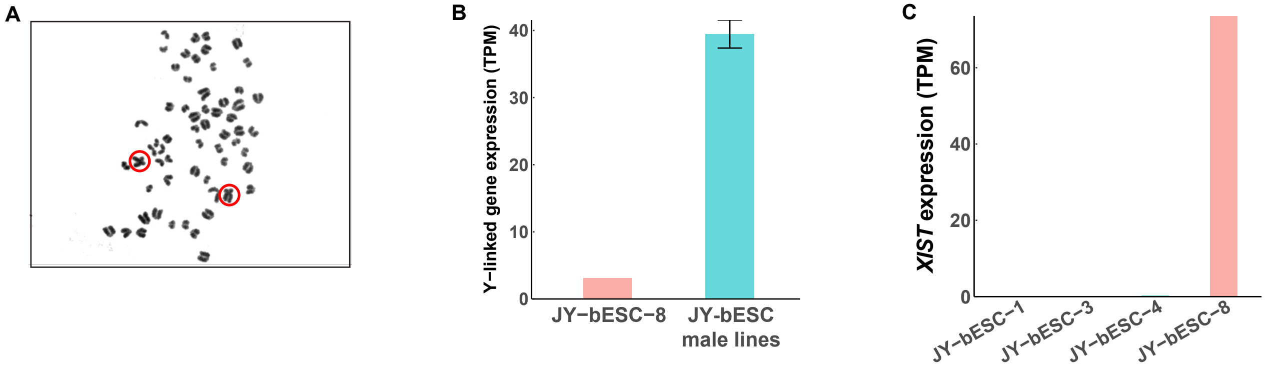

### Suplemental figure 7

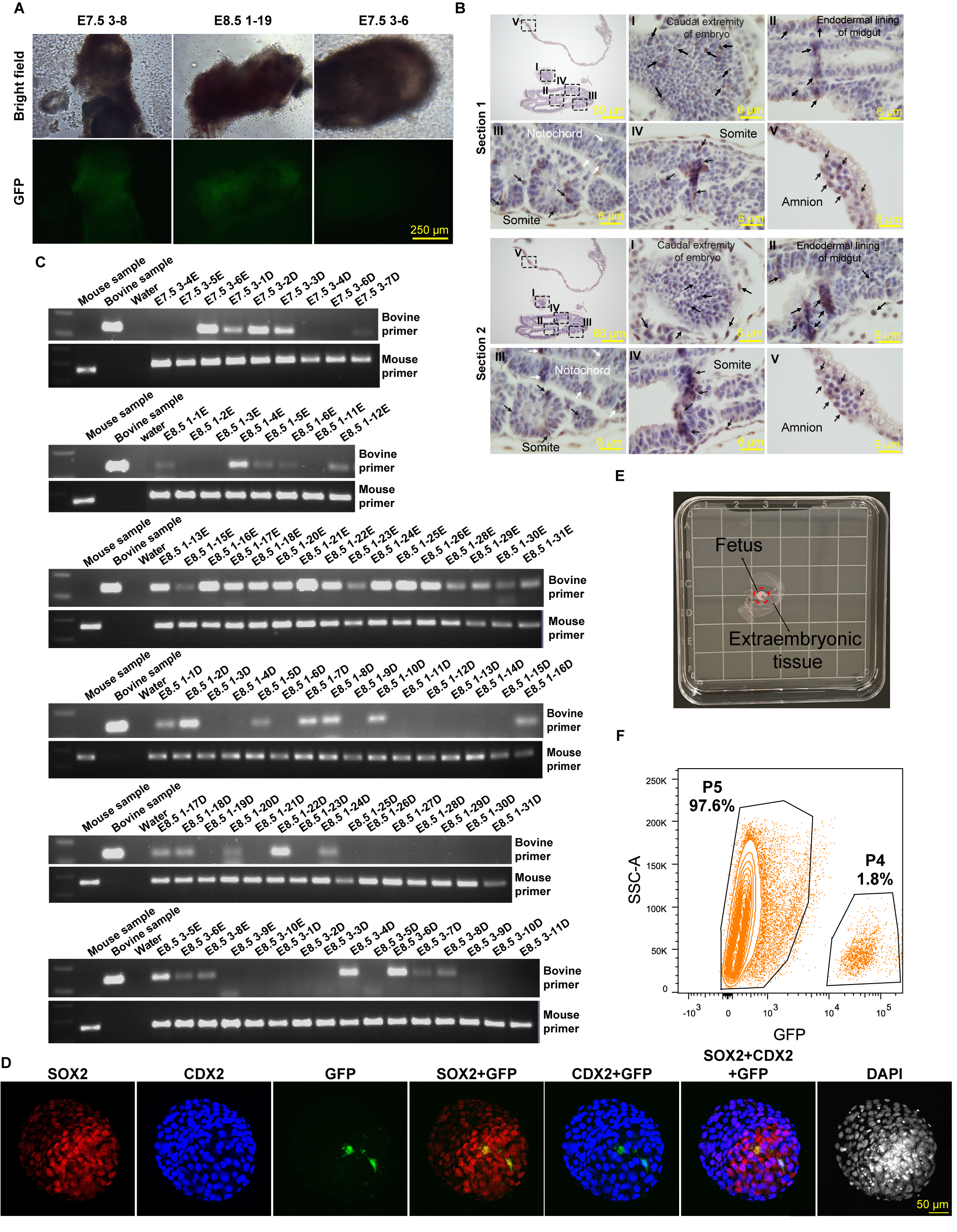

### Suplemental figure 8

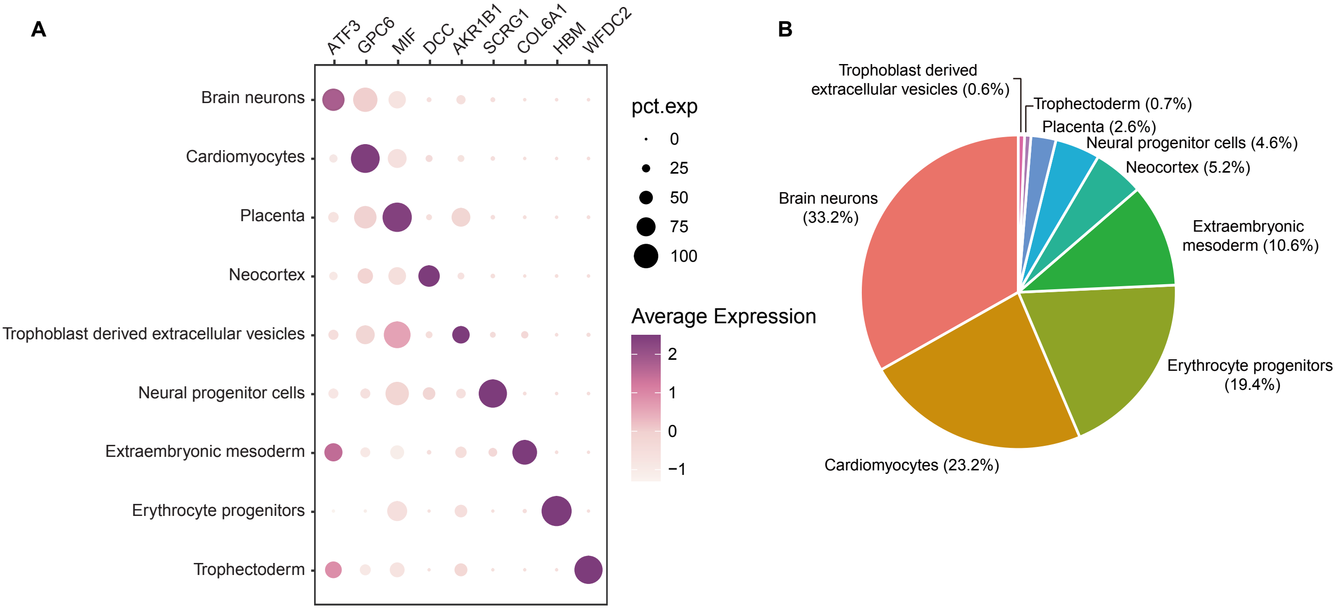
