## Supplementary material for "Bovine Formative Embryonic Stem Cell Plasticity in Embryonic and Extraembryonic Differentiation": Suplemental information

**Figure. S1.** **Derivation and Characterization of bESCs.** (*A*) Relative expression of pluripotency genes by biPSCs in TiF and TiF2 medium. Mean ± sd, n = 3, biological replicates. (*B*) The morphology of biPSCs in TiF2 medium. (*C*) Flow cytometry analysis of CRIPSR-Cas9 edited JY-bESCs with GFP and frame-corrected Cas9 reporter (top right). (*D*) JY-bESC GFP+ line with Cas9 out-of-frame reporter after CRISPR-Cas9 editing. (*E*) F-actin staining of GFP-positive JY-bESCs cultured on Matrigel for 5 days. (*F*) EBs formed by JY-bESCs. (*G*) Negative control immunostaining of JY-bESCs with only secondary antibodies. (*H*) Negative control immunostaining of EBs with only secondary antibodies. (*I*) A representative teratoma formed by JY-bESCs. (*J*) Inner layer of the Z-stack scanning for immunostaining images of PGCLCs for TFAP2C, PRDM1 and SOX17 four days after induction (Z = 18). (*K*) Quantification of TFAP2C, PRDM1, and SOX17 triple-positive cells from three lines of bPGCLC based on Z-stack scanning of immunostaining images. Mean ± sd, n = 3, biological replicates.

**Figure. S2. Comparative Transcriptomic Analysis of PSCs.** (*A*) PCA comparison of JY-bESCs to previously published bovine embryos (ICM, APE, ERSE [GSE53387] and blastocyst [BL; GSE52415 and GSE110036]). (*B*) Heatmap of pluripotency markers across four mouse cell lines: mESC_Yang (mouse naive ESC [GSE89303]), mpESC_Liu (mouse primed ESC [GSE92635]), mFormative_Kinoshita (mouse formative ESC [GSE131556]), and mEPSC_Yang (mouse extended ESC [GSE89303]). (*C*) Heatmap of pluripotency markers across four human cell lines: hnESC_Bi (human naive ESC [GSE174771]), hpESC_Bi (human primed ESC [GSE174771]), hFormative_Yu (human formative [GSE135989]), and hEPSC_Yang (human extended ESC [GSE89303]).

**Figure. S3.** **Comparative Transcriptomic Analysis of PSCs.** (*A*) Heatmap of epigenetic modification-related genes across different bPSCs (bpESC [GSE110036], bEPSC [GSE129760] and biPSC [GSE169624]). (*B*) Heatmap comparison of DEGs between JY-bESCs and bMSCs (GSE180931).

**Figure. S4.** **Comparative Transcriptomic Analysis of PSCs.** (*A*) PCA projection of eY-bESCs' pluripotency onto that of mouse PSCs. (*B*) Heatmap comparison of JY-bESCs, hFormatives_Kioshita (GSE131551), and mFormatives_Kioshita (GSE131556). Key genes related to pluripotency are listed on the left. Significantly enriched GO:BP terms are displayed on the right. (*C*) Heatmap comparison of JY-bESCs, hpESC_Bi (GSE174771), and mpESC_Liu (GSE92635). Key genes related to pluripotency are listed on the left. Significantly enriched GO:BP terms are displayed on the right. (*D)* Venn diagram showing the overlap of unique genes in JY-bESCs and bovine embryos. JY-bESCs unique genes were identified through a comparative analysis of JY-bESCs, hFormatives_Kinoshita, and mFormatives_Kinoshita. Bovine embryo unique genes were identified by comparing previously published RNA-seq data of bovine, human, and mouse embryos. Differential gene expression analysis was performed between each pair within their respective three groups, identifying genes that were highly expressed exclusively in JY-bESCs or bovine embryos as unique genes.

**Figure. S5. Species-specific Metabolic Pathway Analysis.** (*A*) Bar plot of KEGG pathways enriched by the top 2,000 genes primarily contributing to the variance of PC3 in the 3D PCA. (*B-F*) Boxplot of Log2(TPM+1) expression levels for genes involved in various KEGG pathways across different species (cattle, human, and mouse), identified via KEGG analysis of the top 500 genes primarily contributing to PC3 in PCA. (*B*) Carbon Metabolism; (*C*) N-glycan Biosynthesis, (*D*) Reactive Oxygen Species Processes, (*E*) Biosynthesis of Cofactors, and (*F*) Purine Synthesis. Cattle (n = 31), human (n = 19), and mouse (n = 25). (*G*) Boxplot of Log2(TPM+1) expression levels for genes involved in glycerolipids across different species (cattle, human, and mouse). (*H*) Barplot of Log2(TPM+1) expression levels of *TPI1* and *GPD1* in different bovine embryos and PSCs.

**Figure. S6.** **X Chromosome Inactivation (XCI) Analysis of PSCs.** (*A*) Karyotypes of JY-bESC-8 displaying two X chromosomes. (*B*) Comparison of the sum of five Y-linked genes (*OFD1Y*, *USP9Y*, *ZRSR2Y*, *DDX3Y*, and *EIF2S3Y*) between a female line (JY-bESC-8) and three male lines (JY-bESC-1, JY-bESC-3, JY-bESC-4). (*C*) Expression of the *XIST* gene in four JY-bESC lines.

**Figure. S7.** **Inter- and Intra-species Chimeric Embryos Generated from JY-bESCs.** (*A***)** GFP positive (first two panel) and negative (the third panel) E7.5 and E8.5 mouse-bovine chimeric embryos generated with GFP positive bESCs. (*B*) Serial IHC staining images of E7.5 mouse-bovine chimeric embryo using the GFP antibody. The arrows point to the positive regions. (*C*) PCR analysis of JY-bESCs' contribution in E7.5 and 8.5 mouse decidual tissues and embryo proper using bovine- or mouse-specific primers. (*D*) Additional images of whole-mount immunofluorescence of bovine chimeric blastocysts. (*E*) A flushed Day 25 bovine chimeric embryo showing the fetus and extraembryonic tissue. Each square = 16×16 mm. (*F*) Flow cytometry analysis of GFP positive (P4) and negative (P5) cells in Day 25 *in vivo* developed bovine chimeric fetuses.

**Figure. S8.** **10x scRNA-seq Analysis.** (*A*) Dot plot of expression levels of representative markers in each cluster. (*B*) Pie chart of the proportions of different cell types in GFP+ cells of two Day 25 bovine chimeric embryos.
